# Rhythmic coordination of hippocampal-prefrontal ensembles for odor-place associative memory and decision making

**DOI:** 10.1101/2020.06.08.140939

**Authors:** Claire A. Symanski, John H. Bladon, Emi T. Kullberg, Paul Miller, Shantanu P. Jadhav

## Abstract

Memory-guided decision making involves long-range coordination across sensory and cognitive brain networks, with key roles for the hippocampus and prefrontal cortex (PFC). To investigate these coordination mechanisms, we monitored activity in hippocampus (CA1), PFC, and olfactory bulb in rats performing an odor-place associative memory guided decision task on a T-maze. During odor sampling, the beta (20-30 Hz) and respiratory (7-8 Hz) rhythms (RR) were prominent across the three regions, with CA1-PFC beta and RR coherence enhanced during the odor-cued decision making period. Beta phase modulation of CA1 and PFC neuronal activity during this period was linked to accurate decisions, suggesting that this temporal modulation influences sensory-cued decision making. Single neurons and ensembles in both CA1 and PFC encoded and predicted animals’ upcoming choices. Our findings indicate that rhythmic coordination within the hippocampal-prefrontal network supports utilization of odor cues for memory-guided decision making.

## INTRODUCTION

The ability to recall associations from memory and use them to guide behavior is a key aspect of cognition across species. Animals can associate sensory cues in the environment with rewarding and noxious experiences and utilize these cues for adaptive behavior. Memory-guided decision making demonstrates the brain’s remarkable ability to link familiar cues with actions and beneficial outcomes, but little is known about the mechanisms responsible for this cognitive function.

Neurons that encode learned associations and reflect upcoming choice behavior have been reported in multiple regions in different sensory modalities (Allen et al., 2016; Fitzgerald et al., 2011; Harvey et al., 2012; Igarashi et al., 2014; Johnson and Redish, 2007; McKenzie et al., 2014; Moita et al., 2003; Otto and Eichenbaum, 1992b; Schoenbaum and Eichenbaum, 1995b; Shadlen and Newsome, 2001; Singer et al., 2013; Wirth et al., 2009; Wirth et al., 2003; Yanike et al., 2004). However, sensory cued decision-making based on learned associations necessarily involves a brain-wide network that links primary sensory areas, the medial temporal lobe, and higher cortical areas involved in executive function. Numerous studies have highlighted the significance of the hippocampus and prefrontal cortex (PFC) in cognitive processing related to memory and decision making (Battaglia et al., 2011; Euston et al., 2012; Floresco et al., 1997; Lee and Solivan, 2008; Miller and Cohen, 2001). Both regions are known to encode behaviorally relevant cues and task features (Gothard et al., 1996; Hyman et al., 2012; Wiener et al., 1989; Wirth et al., 2009; Wirth et al., 2003; Yanike et al., 2004), and have been shown to play key roles in memory recall (Fortin et al., 2004; Hasegawa, 2000; Siegle and Wilson, 2014; Wiltgen et al., 2004). Notably, coordinated activity between the hippocampus and PFC, supported by bidirectional anatomical connections (Cenquizca and Swanson, 2007; Delatour and Witter, 2002; Ito et al., 2015), has been shown to be critical for learning and memory-guided behavior (Maharjan et al., 2018; Place et al., 2016; Shin and Jadhav, 2016; Shin et al., 2019; Yu and Frank, 2015; Zielinski et al., 2020). Therefore, we focused on the coordinated interactions between hippocampus and PFC as a potential key mechanism through which learned associations are recalled and translated into memory-guided decisions.

Several studies have shown that rhythmic network oscillations in the local field potential (LFP) are involved in long-range interactions between the hippocampus and PFC (Benchenane et al., 2011; Buzsaki and Draguhn, 2004; Colgin, 2011; Gordon, 2011; Jones and Wilson, 2005; Shin and Jadhav, 2016). In particular, phase coherence in distinct frequency bands across this network has been suggested as a mechanism for network coordination underlying mnemonic functions. Notably, hippocampal-prefrontal coherence in the theta rhythm (6-12 Hz) and phase-locked spiking plays a role in spatial working memory and acquisition of spatial tasks (Benchenane et al., 2010; Gordon, 2011; Hyman et al., 2010; Jones and Wilson, 2005). However, whether similar mechanisms of coordination between hippocampus and PFC underlie decision making based on sensory cued associations is unclear.

Rodents rely heavily on odor cues for navigation and foraging, so odor memories are highly salient and robust (Abraham et al., 2004; Eichenbaum, 1998; Rinberg et al., 2006; Uchida and Mainen, 2003), making olfactory memory tasks ideally suited for studying memory-guided decision making. Previous studies using odor memory tasks have found prominent beta (20-30 Hz) oscillations in olfactory regions and the medial temporal lobe during cue sampling, suggesting that the beta rhythm acts as a potential mode of long-range communication for olfactory information processing (Frederick et al., 2016; Igarashi et al., 2014; Kay and Beshel, 2010; Rangel et al., 2016; Stopfer et al., 2003). In addition to the beta rhythm, the respiratory rhythm (RR; 7-8 Hz), driven by the animal’s breathing cycle, is also prominent in the hippocampus during mnemonic processing of odor stimuli (Karalis and Sirota, 2022; Kay, 2005; Kepecs et al., 2006; Lockmann et al., 2016; Nguyen Chi et al., 2016; Verhagen et al., 2007). However, not much is known about the roles of these rhythms in coordinating activity in the hippocampal-prefrontal network during odor-cued decision making.

To elucidate these mechanisms, we employed an odor-place association task in which rats were required to choose the correct trajectory on a T-maze by recalling and utilizing familiar associations between odor cues and reward locations. While rats were performing the task, we recorded simultaneously from the hippocampus and PFC. Given the involvement of both of these regions in odor-memory tasks (Alvarez et al., 2002; Eichenbaum et al., 1986; Fujisawa et al., 2008; Fujisawa and Buzsaki, 2011; Martin et al., 2007; Otto and Eichenbaum, 1992a; Peters et al., 2013; Place et al., 2016), and the role of hippocampal-prefrontal networks in memory-guided behavior (Churchwell et al., 2010; Fortin et al., 2004; Hasegawa, 2000; Moser and Moser, 1998; Wiltgen et al., 2004; Wiltgen et al., 2010), we hypothesized that rhythmic activity in this network may govern the cellular representation of odor-cued decisions underlying behavioral choices. We also monitored LFP activity in the olfactory bulb (OB) in addition to CA1 and PFC, for robust samplings of previously reported olfactory rhythms such as beta and RR, which have been implicated in cognitive processing of olfactory stimuli (Kay, 2014; Kay et al., 2009). Our results point to a role of beta and RR rhythms in coordinating the olfactory-hippocampal-prefrontal network for utilizing learned odor-place associations to inform decisions, and shed light on the cellular and network mechanisms underlying this process.

## RESULTS

### Odor-place associative memory and decision-making task

The odor-cued spatial associative memory task required rats to sniff at an odor port where one of two possible odors was presented using a calibrated olfactometer (see **Methods**). The rats were required to choose the correct associated reward arm on the T-maze based on the sampled odor identity, where they would receive a reward of evaporated milk (**Fig. 1a**).

**Figure 1:**
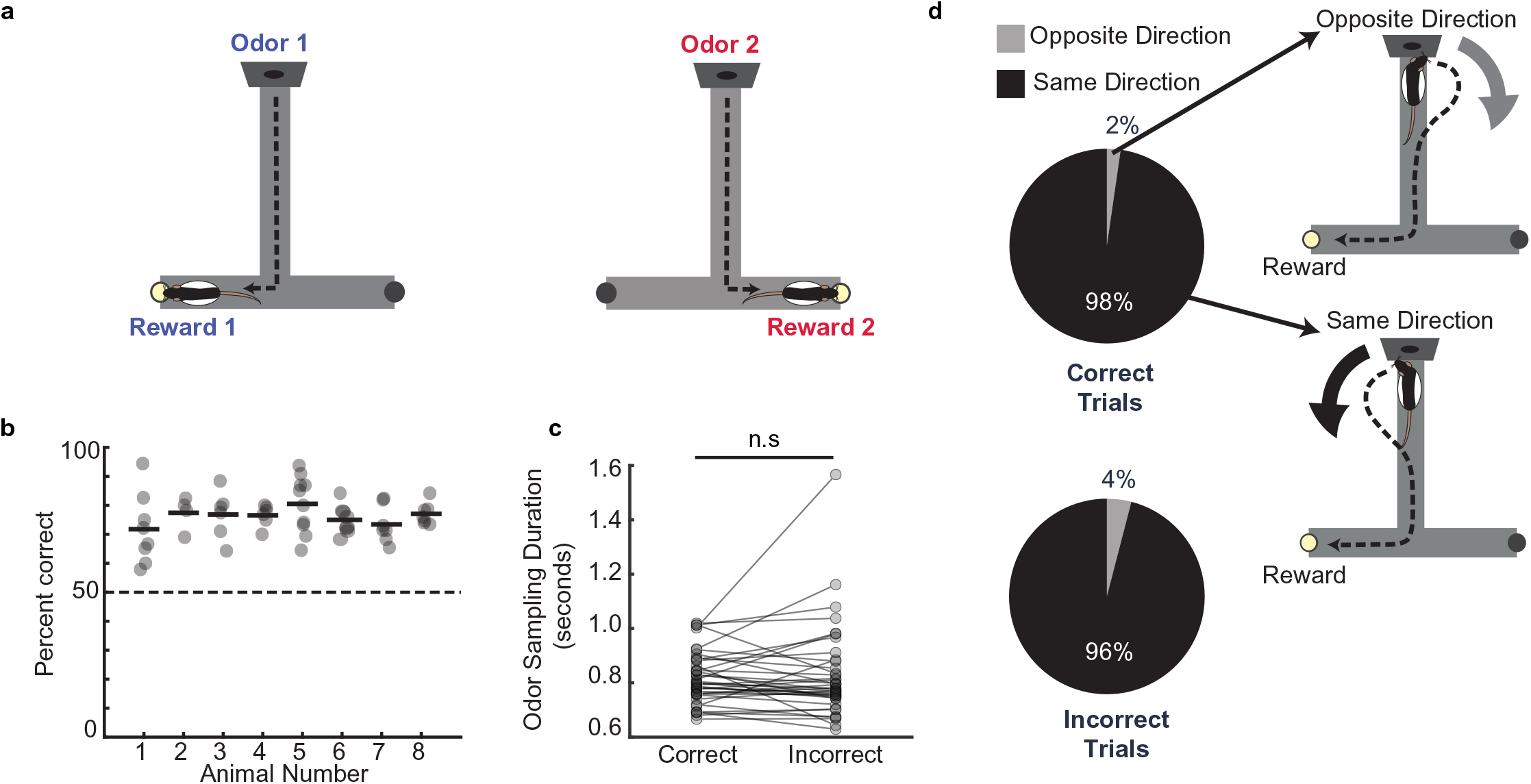
Odor–place associative memory and decision-making task. a. Schematic of the odor-cued T-maze task. Odors 1 (heptanol) and 2 (ethyl butyrate) were delivered at the odor port in pseudo-random order. Presentation of Odors 1 and 2 was associated with milk reward at Reward locations 1 and 2, respectively. Animals had to recall odor-space associations on each trial and utilize the association to choose the correct reward location. b. Performance of each animal (*n* = 8) on the odor-cued T-maze task (animals 1 - 5) or truncated odor-cued task (animals 6 - 8; truncated task, without spatial delay period) across multiple sessions (grey dots). Animal average is indicated by black bars. Dashed line indicates chance level. c. Odor sampling duration across all sessions (n = 38) on correct and incorrect trials (signed-rank test, p = 0.71). d. Turn direction away from odor port in relation to chosen reward well. Pie charts indicate the fraction of trials in which, at the odor port, animals turned in the same direction versus the opposite direction as the reward well that they would ultimately choose. Sessions in which animals ran the truncated task without spatial delay period were excluded. (correct trials: n = 1624 same direction, 32 opposite direction, binomial test, p = 0; incorrect trials: n = 499 same direction, 21 opposite direction, p = 1.4e-108).

Rats were habituated to the maze and pre-trained on the task before surgical implant of a tetrode microdrive array for recording neural data. Following post-operative recovery and during electrophysiological recording, animals maintained a high level of performance on the task, indicating accurate decision making based on cued recall of odor-place associations (**Fig. 1b**, *n* = 8 rats, 77.0% ± 1.3%, mean ± s.e.m.). Rats were required to hold their nose in the odor port for a minimum of 0.50 seconds on each trial, but could continue sniffing the odor for any length of time after the minimum threshold was reached. The odor was continuously dispensed for the entire duration of time that the rat held its nose in the odor port and was only turned off once the rat disengaged from the odor-port. The average odor-sampling duration before odor port disengagement and odor offset was 0.82 ± 0.02 seconds (mean ± s.e.m. across sessions), and this duration was similar between correct and incorrect trials (**Fig. 1c**; signed-rank test, *p* = 0.71, distribution for all trials shown in **Supplementary Fig. 1a**). The animals’ average velocity during the task showed a decrease in speed from the pre-odor period to the odor sampling period in the odor port, followed by an increase in speed after they left the odor port to run to the reward location (**Supplementary Fig. 1b-c**). We observed rapid movement away from the odor-port after odor-port disengagement (**Supplementary Fig. 1d**). In two animals, a thermocouple was implanted in the nasal cavity to measure the sniff rhythm (see **Methods**). There was a small but significant increase in sniff rate during the odor sampling period (7.1 ± 0.39 Hz, mean ± s.e.m.) compared to time matched pre-odor periods (6.2 ± 0.29 Hz, mean ± s.e.m.) (**Supplementary Fig. 1e**).

Notably, we observed that on a majority of trials (95.4% ± 0.12%), the animals’ turn direction away from the odor port matched the direction of the T-maze reward arm that they would ultimately choose on that trial. This behavioral phenomenon was not required for successful performance of the task, and it occurred regardless of whether the trial was correct or incorrect (**Fig. 1d**; binomial tests, correct trials: *p* = 0; incorrect trials: p = 1.4e-108). This observation indicates that the rats recall the odor-place association and choose the reward location for each trial *during* the odor sampling period, before exiting the odor port to run toward the reward. The time of disengagement from the odor port thus provides a trial-by-trial estimate of the moment at which the animal executes the decision. The odor sampling period corresponds to odor-cued recall of the learned association and priming of the subsequent decision to turn toward the reward location, with a behavioral report of the decision occurring at odor port offset. We therefore termed this odor sampling period as “the decision-making period,” since it provides a temporal window between odor onset and odor port exit to investigate mechanisms underlying odor-cued decision making.

### Beta and RR coherence is elevated during odor sampling and decision making

We therefore focused on the decision-making period and first sought to determine the network dynamics that underlie coordination of brain regions during this period. We used a tetrode microdrive array to record local field potentials (LFPs) and single units from the dorsal CA1 region of the hippocampus, the prelimbic region of the prefrontal cortex (PFC, primarily prelimbic area), and the olfactory bulb (OB, only LFP) in rats as they performed the odor-place association task (see **Methods**; **Supplementary Fig. 2a-b**). The thermocouple signal and LFP traces from CA1, PFC, and OB from an example trial are shown in **Fig. 2a**, along with the same LFP signals filtered in the 20-30 Hz band (additional example shown in **Supplementary Fig. 2c**).

**Figure 2:**
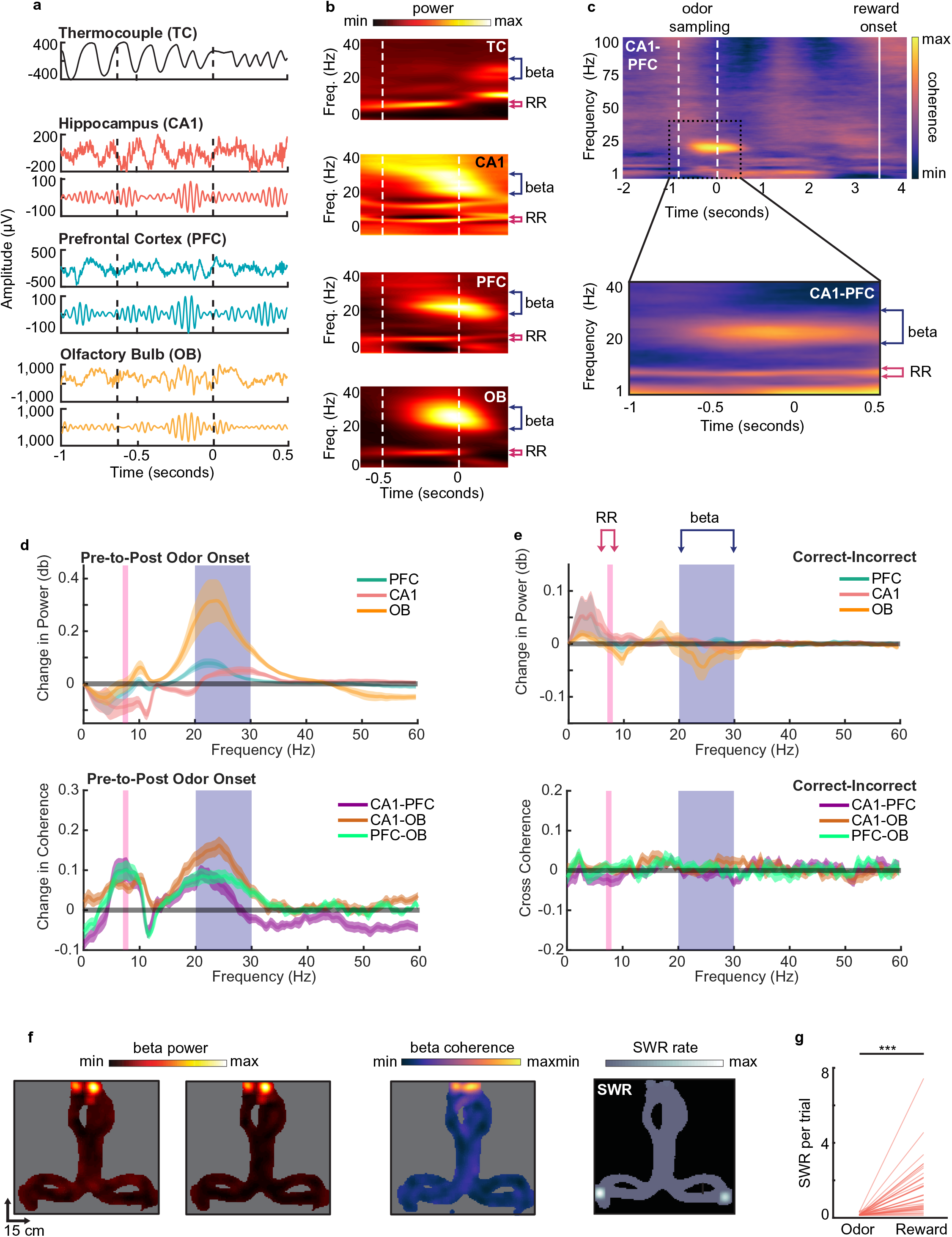
Beta rhythms support decision making based on odor-place associations. a. Examples of thermocouple and LFP traces from one tetrode in each region during presentation of odor from one trial, aligned to odor port disengagement. Area between dashed lines indicates odor sampling period. Top to bottom: Respiratory rhythm recorded via thermocouple, CA1 signal, beta band (20-30 Hz) filtered CA1 signal, PFC signal, beta band filtered PFC signal, OB signal, beta band filtered OB signal. b. Time-frequency plot showing power spectra aligned to odor offset. Color scale represents z-scored power. Area between dashed lines indicates average odor sampling period. Beta band is marked by blue bracket (20-30 Hz). RR band is marked by pink bracket (7-8 Hz). Thermocouple signal (TC), n = 12 sessions, max 0.69, min −0.31; CA1: n = 26 sessions, max 0.26, min −0.35; PFC: n = 26 sessions, max 0.60, min −0.16; and OB: n = 26 sessions, max 2.17, min −0.24. c. Top: CA1-PFC coherence spectra for one animal during the full task time window from odor sampling (area between white dashed lines, aligned to odor offset) to average reward onset time (solid line) (n = 4 sessions, max 0.84; min −0.41). Bottom: CA1-PFC coherence aligned to odor offset across all animals (n = 38 sessions, max 0.51; min −0.34). Color scale represents z-scored power. d. Change in PFC, CA1 and OB LFP power(top) and coherence(bottom) from pre odor period to odor period. Beta band shows significant increase in power (signed-rank tests, n = 26 sessions, CA1 p=2.84e-3, PFC p=1.80e-4, OB p=8.3e-6), while both beta and RR bands show significant increase in coherence (signed-rank test, n=26 sessions, beta: CA1-PFC p=2.84e-3, CA1-OB p=5.1e-5, PFC-OB p=3.67e-5; RR: CA1-PFC p=3.94e-3, CA1-OB p=6.35e-4, PFC-OB p=2.43e-4) e. Change in PFC, CA1 and OB LFP power(top) and coherence(bottom) from incorrect trials to correct trials. No significant changes were detected (signed rank test, n = 26 sessions). f. Maps of T-maze showing the spatial distribution of z-scored CA1 beta power (left. max 1.49; min −0.19), PFC beta power (middle-left, max 1.63; min −0.18), CA1-PFC beta coherence (middle-right, max 1.16; min −0.19), and SWR rate (right, max 1.34; min 0). Plots are for a single session from one animal. g. Number of SWR events per trial during odor sampling and reward consumption on correct trials (signed-rank test, n = 38 sessions, p =1.1e-7***).

We observed a strong increase in power in the beta band (20-30 Hz) during this decision-making period compared to a time-matched pre-odor period across all three regions (**Fig. 2b,d,** signrank test, CA1 p=2.84e-3, PFC p=1.80e-4, OB p=8.3e-6). Similar increases in beta power during odor discrimination tasks have been reported previously in OB, CA1, and lateral entorhinal cortex (Frederick et al., 2016; Igarashi et al., 2014; Kay and Beshel, 2010; Rangel et al., 2016). The respiratory rhythm (RR, 7-8 Hz) was also prominent in the LFP in all three regions, but did not change significantly following odor onset (**Fig. 2d**). This rhythm, which corresponds to the respiration rate during odor sampling, has previously been shown to be physiologically and mechanistically distinct from the 6-12 Hz hippocampal theta rhythm (Lockmann et al., 2016; Nguyen Chi et al., 2016), although there is overlap between the two frequency bands. Following odor port disengagement and the initiation of running down the track, we observed a small shift in the dominant LFP frequency from RR to the theta band in CA1 and PFC, reflecting the change in behavioral state (**Supplementary Fig. 2e**).

Odor sampling during the decision-making period thus drives prominent increases in beta rhythm power in OB, CA1 and PFC. During the decision-making period, we also found phase-amplitude coupling between beta and RR rhythms in all three regions, consistent with previous findings (Lockmann et al., 2016) (**Supplemental Fig. 2f),** and suggesting a mechanistic relationship between the two rhythms. Despite this relationship, previous literature shows that the two rhythms are differentially generated and maintained in the olfactory system and hippocampus and appear to serve different functions in olfactory processing (Kay, 2014; Kay et al., 2009; Martin et al., 2006; Neville and Haberly, 2003), indicating that the beta rhythm is not merely a harmonic of RR. Indeed, power increases were seen during the odor sampling period only in the beta band (**Fig. 2d** signed-rank tests, n = 26 sessions, CA1 p=2.84e-3, PFC p=1.80e-4, OB p=8.3e-6) and not in the RR band (**Fig. 2d** signed-rank tests, n = 26 sessions, CA1 [decrease in power] p=3.94e-3, PFC p=0.551, OB p=0.949).

The hippocampal-prefrontal (CA1-PFC) network exhibited prominent and consistent coherence at the beta frequency band during odor sampling, prior to the decision execution at odor-port offset (**Fig. 2c,** similar figures for CA1-OB and PFC-OB in **Supplementary Fig. 3b & c**). This coherence was specific to the odor-sampling period: it significantly increased compared to time-matched pre-odor periods (**Fig 2d** bottom, signed-rank test, CA1-OB p = 5.1e-5, CA1-PFC p = 2.8e-3, PFC-OB p = 3.6e-5), and diminished shortly after the rat exited the odor port, once the decision had been made (**Fig. 2c**). CA1-PFC coherence in the RR band also increased from the pre-odor to the odor-sampling period (**Fig. 2d,** signed-rank test, CA1-OB p = 6.4e-4, CA1-PFC p = 3.9e-3, PFC-OB p = 2.4e-4). During running, after the decision-making period, the prominent low frequency RR coherence shifted to be slightly higher and centered on the canonical theta band (6-12 Hz), similar to what was observed in the LFP power spectrum (**Supplementary Fig. 3a**). Beta and RR coherence between the CA1-OB and PFC-OB regions also increased during odor sampling periods (**Fig. 2d**), with similar temporal dynamics as CA1-PFC beta coherence.

To control for the possibility that elevated beta coherence during the decision-making period may simply be a reflection of passive movement preparation, as has been observed in sensorimotor cortex (Donoghue et al., 1998), we compared CA1-PFC beta coherence during the last 500 ms of the decision-making period with the 500 ms just prior to reward well exit. We confirmed that beta coherence observed during the decision-making period was significantly higher than coherence at the reward well. **(Supplementary Figure 3f, g,** signed-rank test, CA1-PFC *p* = 0.01, CA1-OB p=0.0012, PFC-OB p=0.0024). Interestingly, this effect in RR was observed for CA1-OB and PFC-OB, but not between CA1-PFC (**Supplementary Figure 3f, g**, signed-rank test CA1-PFC p=0.569, CA1-OB p=0.0044, PFC-OB p=0.00064), suggesting that in contrast to CA1-PFC beta coherence, the increase in CA1-PFC RR coherence was not specific to the decision making period. In an additional control, we also confirmed that this elevated beta coherence was specifically seen during odor-cued decision making and was not present in sessions in which only air was presented at the odor-port, similar to previous reports for beta coherence in CA1-entorhinal cortex (Igarashi et al., 2014) (see **Methods** for description of air-only sessions). In these air-only sessions, reward was given on random trials at the reward locations, even though no odors were presented. RR coherence was unchanged between odor- and air-cued sessions, but beta coherence was significantly lower in the absence of odor cues during air-only sessions (**Supplementary Fig. 3d-e**).

We also examined if these beta rhythm dynamics were limited to the odor sampling location on the maze. Indeed, CA1-PFC network beta power and coherence were clearly isolated to the region near the odor port (**Fig. 2f**). Further, since hippocampal sharp-wave ripples (SWRs) have been proposed as a mechanism for memory retrieval, decision making, and planning in some tasks (Carr et al., 2011; Joo and Frank, 2018; Norman et al., 2019; Singer et al., 2013), we investigated the occurrence of SWRs during our task. SWR events occurred frequently at the two reward locations, as expected (Buzsaki et al., 1983). However, a distinct lack of SWR events at the odor port (**Figs. 2f** and **2g**; signed-rank test, *p* = 1.1e-7) suggests that SWR replay is unlikely to play a direct role in decision making based on recalled associative memories for this well-learned task.

CA1-PFC beta coherence was thus specifically enhanced during the odor sampling and decision making period, suggesting that beta rhythms can play a role in odor-cued decision making in this task. The prevalence of RR rhythms also suggests a possible role in organizing sensory information transfer in odor tasks, as suggested by previous studies (Kay, 2005; Kepecs et al., 2006; Nguyen Chi et al., 2016). We, however, did not find a direct relationship between the strength of either beta coherence or RR coherence to correct performance on the task. There was no significant change in the level of coherence or power in the CA1-PFC network in either frequency band between correct and incorrect trials (**Fig. 2e,** signed-rank tests; all p’s > 0.05). Therefore, while beta coherence in the CA1-PFC network is elevated during odor-cued decision making, the strength of overall oscillatory coordination in the network, as assessed by coherence, may not directly enable a correct decision, suggesting that oscillatory phase modulation of neuronal activity may instead play a role.

### Single neurons in CA1 and PFC exhibit choice selectivity during decision-making period

In addition to LFP, we assessed neuronal activity in the task by recording single units from hippocampal area CA1 (n = 1,309 units) and PFC (n = 717 units) (distribution of neurons across animals shown in **Supplementary Table 1**). Units that were active during run sessions were categorized into pyramidal cells and interneurons based on firing rate and spike width (see **Methods,** CA1: 87% pyramidal, 13% interneurons; PFC: 87% pyramidal, 13% interneurons). Pyramidal cells were then further categorized as task-responsive or task-unresponsive based on whether they exhibited significant changes in firing rate during odor sampling (see **Methods**; n = 125 task-responsive CA1 cells, n = 157 task-responsive PFC cells). We observed an overall increase in spiking during the odor sampling period in the task-responsive cell population, suggesting that the majority of these cells increase their firing rate during this period (**Supplementary Fig. 4a**). Interestingly, however, there was an overall decrease in spiking in the average peri-stimulus time histograms when pooling all spikes from all cells during the odor-sampling period (**Supplementary Fig. 4b**). This suggests a decrease in overall activity in CA1 and PFC networks during odor sampling, with the task-responsive neurons representing a subset of neurons that were especially engaged in the association and decision-making task. Additionally, we confirmed that task responsive neurons were not simply modulated by the animals’ decrease in speed at the odor-port by comparing firing rates during odor sampling to firing rates during reward consumption, when speed was also low. We found task-responsive cells’ firing rates were significantly higher at the odor port than at the reward wells (**Supplementary Fig. 4c**).

A subset of task-responsive cells was selective for specific choices, based on the identity of the odor and the associated choice. Selectivity was calculated by generating a selectivity index (SI), in which the difference between the average firing rate response to each odor on correct trials was divided by the sum of the two responses, giving a value between −1 and 1 (see **Methods**). To determine significance, the SI value for each cell was compared to a distribution of SIs generated by shuffling the odor identities across trials (n = 42 CA1 selective cells, n = 49 PFC selective cells) (**Figs. 3a** and **3b**). Although the number of selective cells that we observed was relatively low, these numbers were consistent with other studies examining single unit responses to odor stimuli in these regions (Otto and Eichenbaum, 1992b; Schoenbaum and Eichenbaum, 1995a; Taxidis et al., 2020). Peri-stimulus time histograms and raster plots aligned to decision time for two example selective cells for CA1 and PFC are shown in **Figs. 3c** and **3g** respectively. Notably, we found that on incorrect trials, selective cells often exhibited responses to the two odors that were opposite to the responses on correct trials (examples shown in **Figs. 3d** and **3h**). We found that this phenomenon held true across the population of selective cells, such that overall, the SIs on correct trials were anti-correlated with the SIs on incorrect trials (**Figs. 3e-f** and **3i-j).** Note that since there are only two possible choices on the task, the behavioral response on an incorrect trial is identical to that of a correct trial of the opposite odor. The fact that the neural responses on trials with identical behavioral responses are similar indicates that selective cells are not simply coding the odor identity but are instead responsive to the animal’s memory-guided decision and upcoming behavioral choice. In contrast, selectivity to the odor identity alone would result in similar responses to a given odor regardless of the upcoming behavior or the ultimate trial outcome. We therefore termed these cells choice-selective cells. This pattern was not significant within the population of neurons that were task-responsive but did not meet the selectivity threshold (**Supplementary Fig. 4d-e**). Histograms showing distribution of absolute selectivity indices between selective and nonselective cells are shown in **Supplementary 4f.**

**Figure 3:**
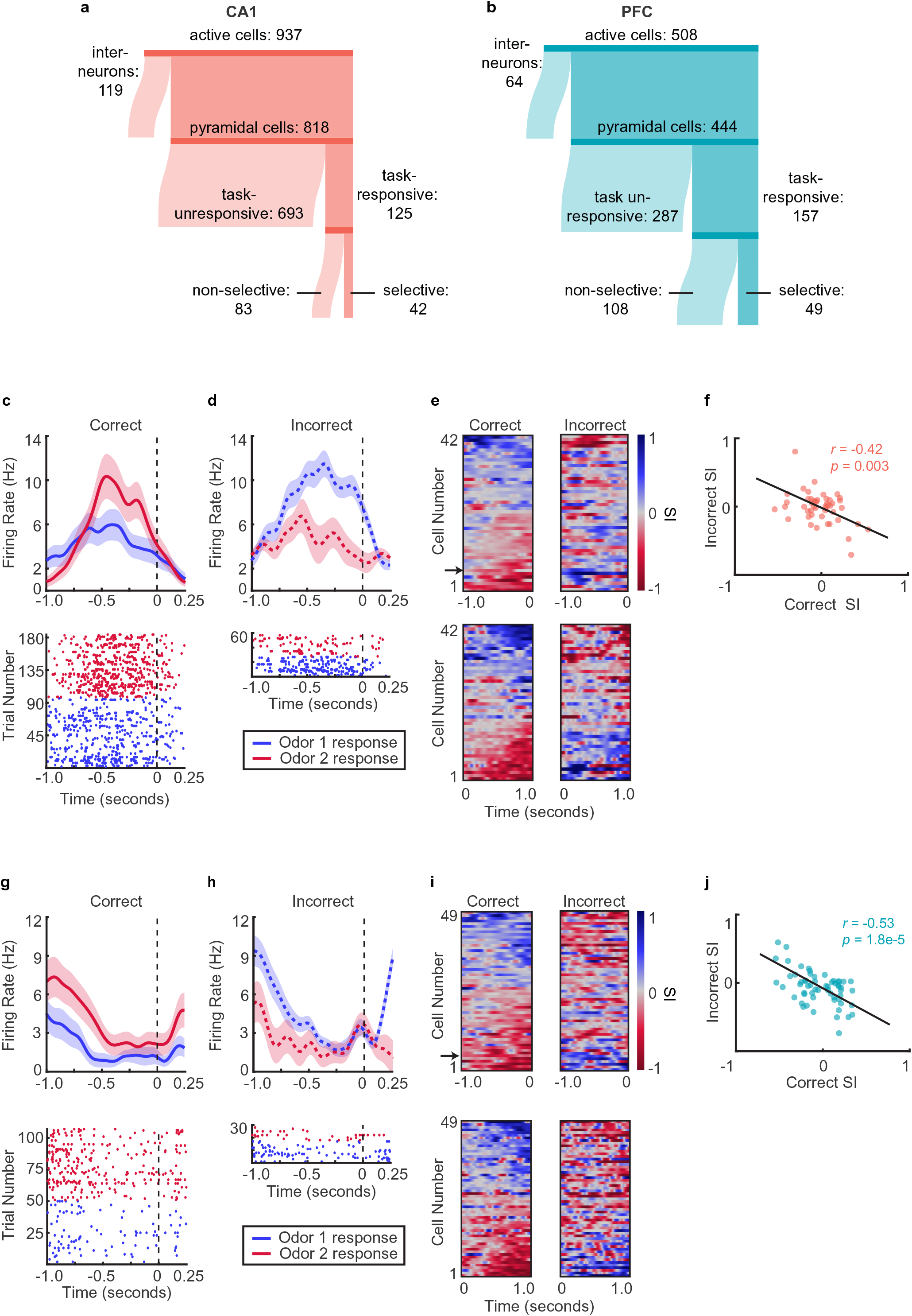
Single neurons in CA1 and PFC exhibit choice-selectivity during decision-making period. a. Sankey diagram showing the number of CA1 cells that were classified into different categories. Sizes of partitions are proportional to raw numbers. b. Same as for a, but for PFC cells. c. Example PSTH and raster plot for a single choice-selective CA1 neuron on correct trials, aligned to odor-port disengagement. Shaded areas indicate s.e.m. Firing rates are shown in Hz, referring to spikes/second. d. Same as for c, but for Incorrect trials. e. Selectivity index (SI) of all choice-selective cells in CA1 on correct trials (left) and incorrect trials (right), aligned to odor-port disengagement. SI is calculated as the difference in firing rate between Odor 1 trials and Odor 2 trials, divided by the sum of these firing rates. SI is color coded, where blue indicates an SI of 1 (absolute Choice 1 preference), red indicates an SI of −1 (absolute Choice 2 preference) and grey indicates an SI of 0 (not selective). Cells are sorted according to peak selectivity on correct trials and sorting order is the same for both plots. f. Correlation between correct trial SI and incorrect trial SI for all choice-selective CA1 cells (n = 42, r = − 0.42, p = 0.003**). g-j. Same as a-e, but for PFC cells. Correlation in j: n = 49, r = −0.53, p = 1.8e-5***.

Putative interneurons were also divided into task-responsive and task-unresponsive, using the same criteria as for putative pyramidal cells (**Fig. 4a**). **Fig. 4b-c** shows interneuron odor selectivity indices and response properties during correct and incorrect trials. For CA1 interneurons, the selectivity response showed anti-correlation during correct versus incorrect trials similar to that of pyramidal cells. PFC interneurons’ SIs on correct and incorrect trials were not anti-correlated, although this could result from the low number of selective PFC interneurons overall.

**Figure 4:**
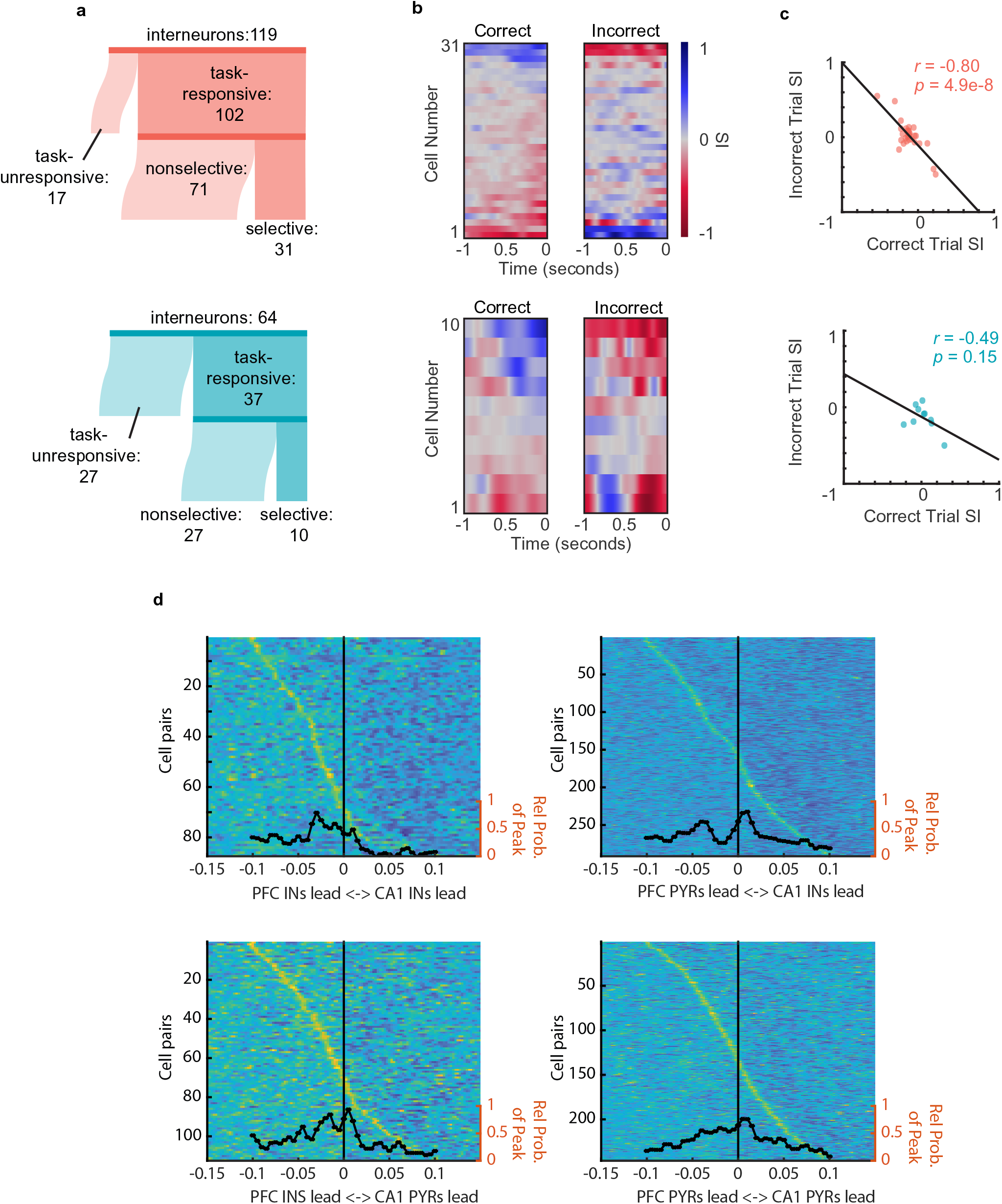
Interneurons exhibit choice-selectivity and temporal coordination during decision-making period. a. Sankey diagram showing the number of CA1 interneurons (top) and PFC interneurons (bottom) that were classified into different categories. Sizes of partitions are proportional to raw numbers. **b.** Selectivity index (SI) of all choice-selective interneurons in CA1 (top) and PFC (bottom) on correct trials (left) and incorrect trials (right), aligned to odor-port disengagement. c. Correlation between correct trial SI and incorrect trial SI for CA1 (n = 31, r = −0.8, p = 4.9e-8) and PFC (n = 10, r = −0.49, p = 0.15) interneurons. d. Histograms and waterfall plots of significantly connected PFC-CA1 cell pairs. Peaks falling above zero indicate CA1 leading, whereas peaks falling below zero indicate PFC leading. One-sample Wilcoxson signed rank tests (H_0_: µ = 0 ms). Top-left: CA1-PFC interneuron pairs, n = 87 pairs, p = 3.5e-8; top-right: CA1 interneuron-PFC pyramidal pairs, n = 283 pairs, p = 2.11e-3; bottom-left: CA1 pyramidal-PFC interneuron pairs, n = 111 pairs, p = 3.4e-3, bottom right: CA1-PFC pyramidal pairs, n = 224 pairs, p = 0.14.

To investigate whether there was evidence of coordination between ensembles in CA1 and PFC, we wanted to determine whether the spiking of neuronal ensembles in the two regions were temporally linked (Jadhav et al., 2016; Siapas et al., 2005). In order to examine this, we computed the normalized spiking cross-correlation for all CA1-PFC pairs of task-responsive neurons during odor sampling (**Fig. 4d, Methods**). Significant peak time lags of cross-correlations were quantified for all CA1-PFC task-responsive putative pyramidal cells (n = 224 pairs), all pairs of task-responsive interneurons (n = 87 pairs), as well as pairs of CA1 interneurons and PFC pyramidal cells (n = 298 pairs), and vice versa (n = 113 pairs). We found a significant skew in CA1-PFC interneuron pairs towards the PFC interneurons leading (sign rank test, x=-.028, p=3.5e-8) (**Fig. 4d**), as well as for CA1 pyramidal cell - PFC interneuron pairs in the same direction (sign rank test, x=-.013, p=0.0034). Additionally, we found a large cluster of CA1 interneuron-PFC pyramidal cell pairs whose cross-correlations peaked around −.035 sec and one at +.01 sec, or approximately the period of a beta cycle.

These results further illustrate temporal relationships during the decision making period, and especially suggest that CA1 interneurons show temporal coordination with PFC task-responsive ensembles in the beta range. The strong skew towards PFC interneuron’s leading CA1 interneurons and CA1 pyramidal cells suggest top down influence of the PFC on CA1 just before the animal begins to execute its decision. Since interneurons have a pre-dominant effect on local network activity, common input to the CA1-PFC network, especially to interneurons, can potentially play a key role in temporally organizing ensemble responses in both regions (Andrianova et al., 2021; Dolleman-van der Weel et al., 2019; Schlecht et al., 2022; Varela et al., 2014).

### CA1 and PFC cells phase lock to beta and respiratory rhythms during decision making

We next asked if there was any relationship between oscillatory phase modulation of neuronal activity and decision accuracy. A subset of cells within the population of task-responsive cells in both CA1 and PFC were phase locked to the beta rhythm during the decision-making period (see **Methods**; CA1: pyr n = 12/86, 14.0%, int n = 29/71, 40.9%; PFC: pyr n = 6/92, 6.5%, int n = 4/33, 12.1%), and many cells in the task-responsive populations were also strongly phase locked to RR (CA1: pyr n = 59/86, 68.6%, int n = 66/71, 92.9%; PFC: pyr n = 15/92, 16.3%, int n = 10/33, 30.3%). Spike-phase histograms for example beta-phase and RR-phase locked cells in CA1 and PFC are shown in **Fig. 5a & Fig 5b** (Rayleigh Z test, CA1: n = 242 spikes, z = 15.4, p = 1.5e-7; PFC: n = 99 spikes, z = 40.9%, p = 0.007), and the preference for all coherent cells is shown in polar plots (**Fig. 5c & d**). Interestingly, even though a higher percentage of cells in both regions were phase locked to RR (**Fig. 5e**, z-test for proportions, CA1: p = 6.8e-23; PFC: p = 8.8e-9) and depth of this modulation was much stronger for RR compared to beta (**Supplementary Fig. 5a**), phase preference across cells was consistent only in CA1, but for both rhythms (**Fig 5c, d,** Rayleigh Z-test, CA1: n=157 beta p=4.3e-6, RR p=2.01e-5; PFC n=125, beta p=0.70, RR p=0.13). Additionally, in both CA1 and PFC, the proportion of cells that were phase locked to both beta *and* RR was no different than chance, given the percentages of cells that were modulated by either rhythm (**Supplementary Fig. 5b**). This suggests that the two rhythms simultaneously modulate different populations of cells, supporting the interpretation that the two rhythms play separate, though cooperative, roles in memory-guided decision making.

**Figure 5:**
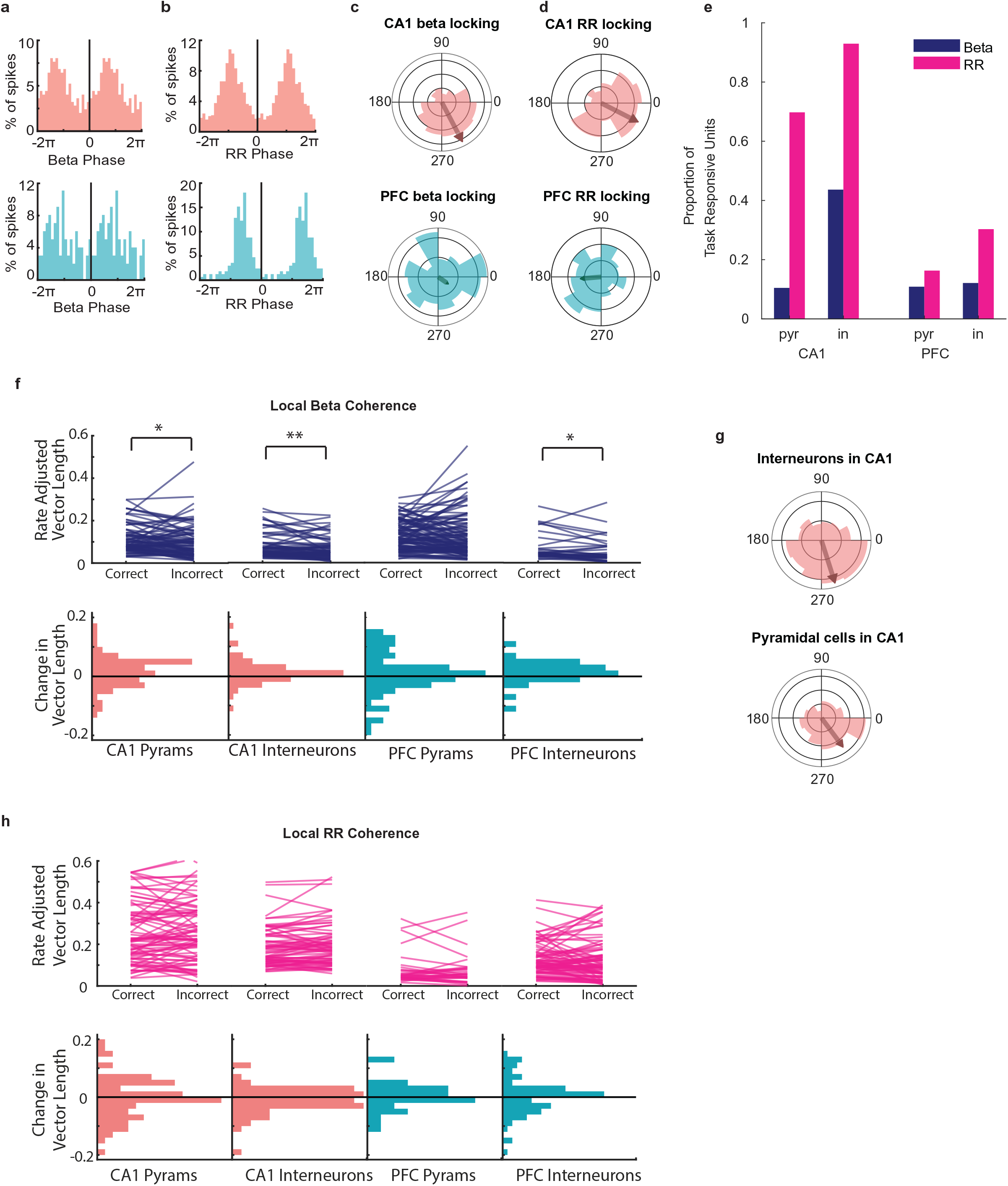
CA1 and PFC cells phase lock to beta and respiratory rhythms during decision making. a. Example spike-phase histograms from two example cells that are phase locked to the beta rhythm. Top: CA1 cell (Rayleigh Z test, n = 242 spikes, z = 15.5, p = 1.5e-7); bottom: PFC cell (n = 99 spikes, z = 4.86, p = 0.007). Phase axes are duplicated for visibility. b. Example spike-phase histograms from two example cells that are phase locked to RR. Top: CA1 cell (Rayleigh Z test, n = 870 spikes, z = 135.3, p = 6.3e-62); bottom: PFC cell (n = 172 spikes, z = 73.9, p = 3.9e-37). c. Polar histogram of preferred beta phases for all task-responsive CA1 (n = 157 cells, Rayleigh Z test, z = 12.22, p = 4.3e-6) and PFC (n = 125 cells, z = 0.36, p = 0.70) cells. Arrows indicate mean phase direction and magnitude. d. Polar histogram of preferred RR phases for all task-responsive CA1 (Rayleigh Z test, z = 8.96, p = 2.01e-5) and PFC cells (z = 2.02, p = 0.13). Arrows indicate mean phase direction and magnitude. e. Percentage of cells in CA1 and PFC that were phase locked to either beta or RR (Beta; CA1: pyr n = 12/86, 14.0%, in n= 29/71, 40.9%; PFC: pyr n = 6/92, 6.5%, in n = 4/33, 12.1%. RR; CA1: pyr n = 59/86, 68.6%, in n=66/71, 92.9%; PFC: pyr n = 15/92, 16.3%, in n=10/33, 30.3%). f. Rate Adjusted Vector Length(top) and histogram of change in vector length from correct to incorrect trials(bottom) for all task-responsive cells in PFC and CA1 relative to local beta rhythm. (signed-rank tests, signed-rank tests, CA1: pyr n = 90, p = 0.019, in n=73, =p2.61e-3; PFC: pyr n = 98, p = 0.25, in n=33, p=0.043, *<.05, **<.01). g. Polar histogram of preferred beta phases for all task responsive CA1 interneurons and pyramidal cells. (pyr n=86, p=2.81e-5, IN n=71, p=0.0179) h. As in (f), but for all cells relative to local RR. (signed-rank tests, no test found significance below p=0.05).

Are cells that are modulated by beta the same population of cells that are selective for the upcoming decision? To test this, we compared the number of pyramidal cells that were both choice-selective *and* phase-locked (within and across regions) to the fraction that would be expected by chance, given the total percentages of cells that are choice-selective or phase-locked. Interestingly, we found that the number of cells that met both criteria was no different than chance for either CA1 or PFC (**Supplementary Fig. 5c**, binomial tests; CA1: p = 0.097; PFC: p = 0.17). We found similar results for interneurons in each region, as well as for RR modulation (**Supplementary Fig. 5d, e**). It should be noted that since such a large majority of neurons in CA1 were significantly phase locked to RR, it is unsurprising that almost all the choice-selective cells were phase locked to this rhythm. However, this result indicates that the choice-selective cells were no more or less likely than non-selective cells to phase lock to RR as well. These results together suggest that the putative neural ensembles that are selective for the primary task parameter of odor-cued associative decisions may not be driven directly by the beta rhythm via prevalence of phase-locking.

We next asked whether the strength of phase locking to the rhythms was still indicative of decision accuracy by comparing correct versus incorrect trials. To do this, we compared each cells mean vector length (MVL) after adjusting for rate differences (Rangel et al., 2016), a measure of non-uniformity in the spike-phase distribution, on correct versus incorrect trials for all task responsive cells. For each cell, we calculated the MVL for the spike-phase distribution on correct trials and incorrect trials separately and compared these two paired distributions. We found interneurons in both CA1 and PFC as well as pyramidal cells in CA1 that were phase-locked to the local beta rhythm exhibited a lower MVL on incorrect trials (**Fig. 5f**, signed-rank tests, CA1: pyr n = 90, p = 0.019, in n=73, =p2.61e-3; PFC: pyr n = 98, p = 0.25, in n=33, p=0.043). There was no significant effect for the RR phase locking (**Fig 5h,** signed-rank test, no significance found), and neither was there for cross-regional beta modulation (**Supplemental Fig 5f**). This suggests that local spiking modulation by the beta rhythm in CA1 and PFC can potentially play a role in supporting accurate utilization of odor-place associations for making decisions. Together with the observed temporal coordination between task-responsive interneurons across regions (**Fig. 4d**), these data suggest that rhythmic coordination between PFC and CA1 via the beta rhythm supports successful memory guided behavior in this paradigm.

The stronger local beta phase modulation in CA1 during correct trials and elevated CA1-PFC beta coherence during decision making periods (**Fig. 2d**) together suggests that beta coordination is related to task performance. The higher beta phase modulation of CA1 and PFC interneurons also suggests that entrainment of interneurons to the network-wide beta rhythm is linked to task performance as well. This also aligns with the finding of temporal relationships in cross-correlations between PFC interneurons and CA1 pyramidal cells and interneurons (**Fig. 4d**). Rhythmic modulation of interneurons, possibly through a common input, can potentially constrain and coordinate network activity, and may play a role in sculpting pyramidal cell ensembles in local circuits to enable processing of odor-cued associations for translation to decisions.

### Neural ensemble responses in CA1 and PFC during decision making predict the upcoming choice

Given the choice-selective responses of single neurons in CA1 and PFC (**Figs. 3-4**), and the relationship of phase modulation to decision accuracy (**Fig. 5**), we next examined how ensemble dynamics underlie the neural representation of decisions informed by odor-cued recall. We considered all cells that were task-responsive, including putative pyramidal cells and interneurons, for the ensemble analyses. For the population of task-responsive cells, we found that the distribution of the peak response times in both CA1 and PFC tiled the entire decision time window (**Fig. 6a**).

**Figure 6:**
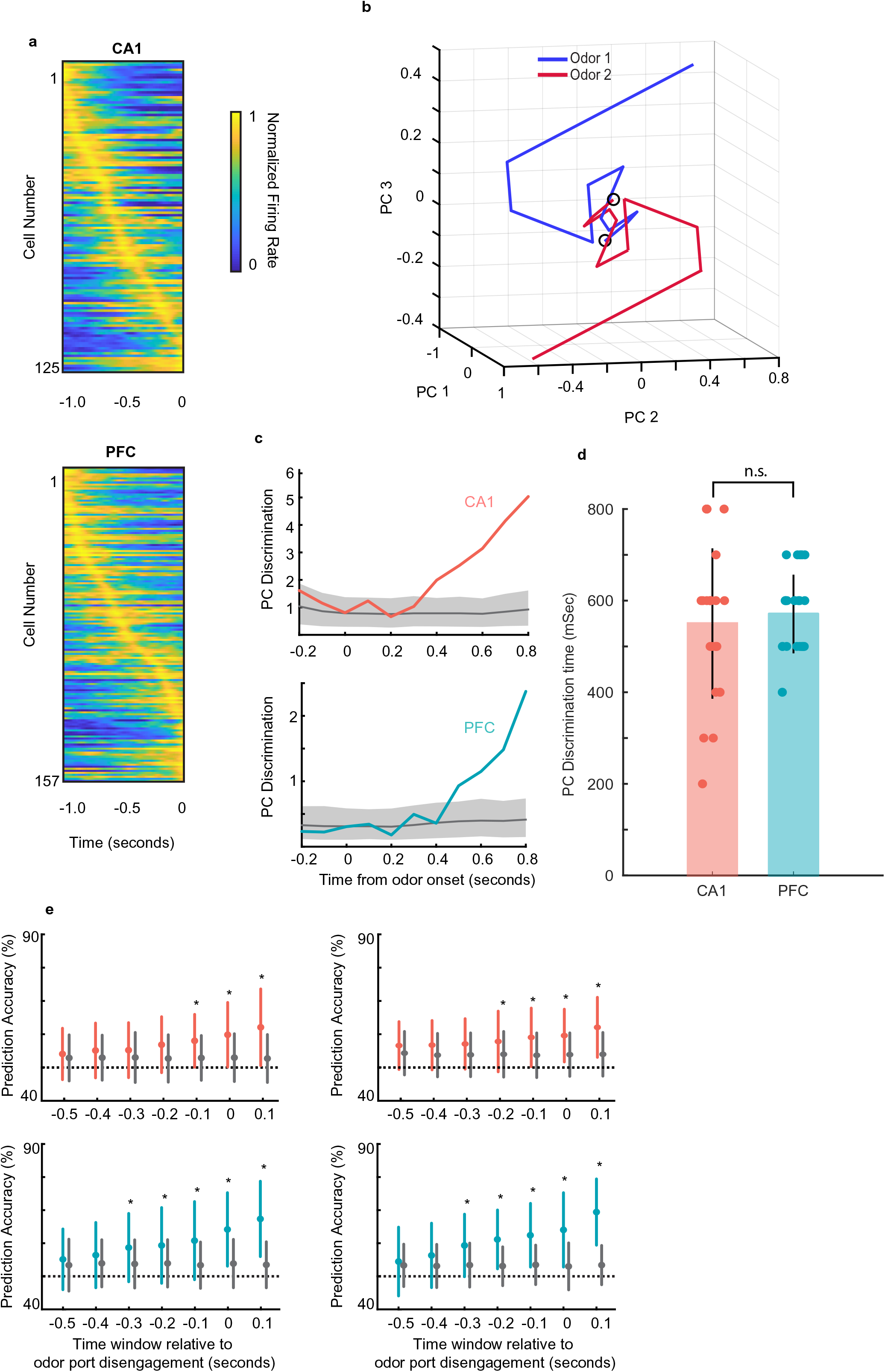
Neural ensemble responses during decision making predict the upcoming choice. a. Normalized firing rate of all task-responsive cells in CA1 (top) and PFC (bottom) during a 1 second window before odor port disengagement. b. Example session with average trajectories of CA1 population activity using the first three PC dimensions for the two choices (n = 43, 45 left-bound (blue) and right-bound (red) trials; n = 12 CA1 neurons in ensemble, 100 ms bins). The two trajectories showing a rapid evolution and separation within a few hundred ms after odor onset. c. Euclidean distance between left-bound and right-bound average PC trajectories for an example CA1 ensemble (top, n = 12 neurons) and PFC ensemble (bottom, n = 10 neurons) was compared to a chance-level distance distribution computed by shuffling the odor identities across trials and creating a null distribution. d. Neural discrimination times for CA1 and PFC locked to odor onset (n = 24 sessions CA1 range: 200-800 ms; PFC range, 400-700 ms) were similar (Fig. 7c; sign rank test, p = 0.68). e. Neural discrimination times during odor sampling aligned to decision time. Colored error bars indicate real data mean ± s.d., whereas gray error bars indicate shuffled data mean ± s.d. Cells used for prediction include all task-responsive putative pyramidal cells in CA1 (top) and PFC (bottom) or all active neurons including interneurons (right). Stars indicate prediction time windows where the fraction of correctly predicted trials was significantly higher than the fraction from the shuffled data (rank-sum tests, * = p < 0.05).

We then examined the temporal evolution of CA1 and PFC ensemble responses in individual sessions to infer the timing of neural discrimination. Principal component analysis (PCA) was applied to visualize the temporal dynamics of population activity during the decision making period, and the Euclidean distance between left-bound and right-bound neural trajectories was used as a measure of neural discrimination (see **Methods**). **Figure 6b** shows an example session with average trajectories of CA1 population activity using the first three PC dimensions for the two choices (n = 12 CA1 neurons in ensemble, 100 ms bins; n = 43 left-bound and n = 45 right-bound trials). The two trajectories show a rapid evolution and separation within a few hundred milliseconds after odor onset. The Euclidean distance between left-bound and right-bound average trajectories was compared to a chance-level distance distribution computed by shuffling the trial identities across trials and creating a null distribution. The timepoint at which the population responses were considered significantly distinct from each other was defined as the first bin at which the real distance between trajectories surpassed the 95% confidence interval of the null (shuffled) distribution (shown in **Fig. 6c** for one example session for CA1 and PFC each. Neural discrimination times for CA1 and PFC (n = 24 sessions across 8 animals; threshold of 4 task-responsive cells, CA1 mean: 200-800 ms; PFC mean: 400-700 ms) were similar (**Fig. 6d**; p = 0.68, sign-rank test).

To determine whether the animal’s upcoming behavior could indeed be predicted by neural ensembles before the decision was executed, we trained a generalized linear model (GLM) to predict the animal’s choice based on ensemble activity during the decision-making period (See **Methods**). We performed this analysis using task-responsive putative pyramidal cells and found that reward choice could be accurately predicted 0.1 seconds prior to odor port disengagement by CA1 ensembles, and 0.3 seconds prior by PFC ensembles (**Fig. 6e**). Additionally, we performed the same analysis, but included both task-responsive pyramidal and interneurons in the ensembles. In this case, we found an improvement in prediction for CA1 ensembles: reward prediction was now accurate starting earlier, at 0.2 seconds prior to decision execution. Prediction by PFC ensembles remained the same. (**Fig. 6e**). To control for the possibility that inclusion of interneurons improved prediction latency for CA1 simply due to the larger number of cells in the training set, we performed this analysis again by resampling the pyramidal cells to match the total number of pyramidal cells plus interneurons. In this control analysis, we found that the choice could again only be predicted at 0.1 seconds prior to the decision execution, the same as what we observed with the original sample of pyramidal cells. These results confirm that the animals utilize the recalled odor-place association and make a spatial choice during this odor-sampling period, which is reflected in the activity of task-responsive neural ensembles.

### Representations of choice and space are maintained independently during delay period

Finally, we investigated whether there was any relationship between the activity of CA1 and PFC ensembles during the odor-cued decision-making period and their spatial activity on the maze as the animals ran through the delay period on the central and side arms toward reward. We examined units from sessions in which animals traversed a long T-maze track (see **Methods**, 26 sessions from 6 rats) in order to assess spatial firing characteristics. A large fraction of cells in CA1 and PFC exhibited spatial activity on the maze (see **Methods** for details on spatial parameters, n = 344 out of 585 CA1 cells; n = 159 out of 288 PFC cells had fields on the track), including units that were both task-responsive (choice selective and non-selective) and task-unresponsive (**Figs. 7a, 7b**; examples of CA1 and PFC choice-selective cells with spatial fields in **Fig. 7a,** examples of CA1 and PFC task-responsive but non-selective cells with spatial fields shown in **Supplementary Fig. 6a**).

**Figure 7:**
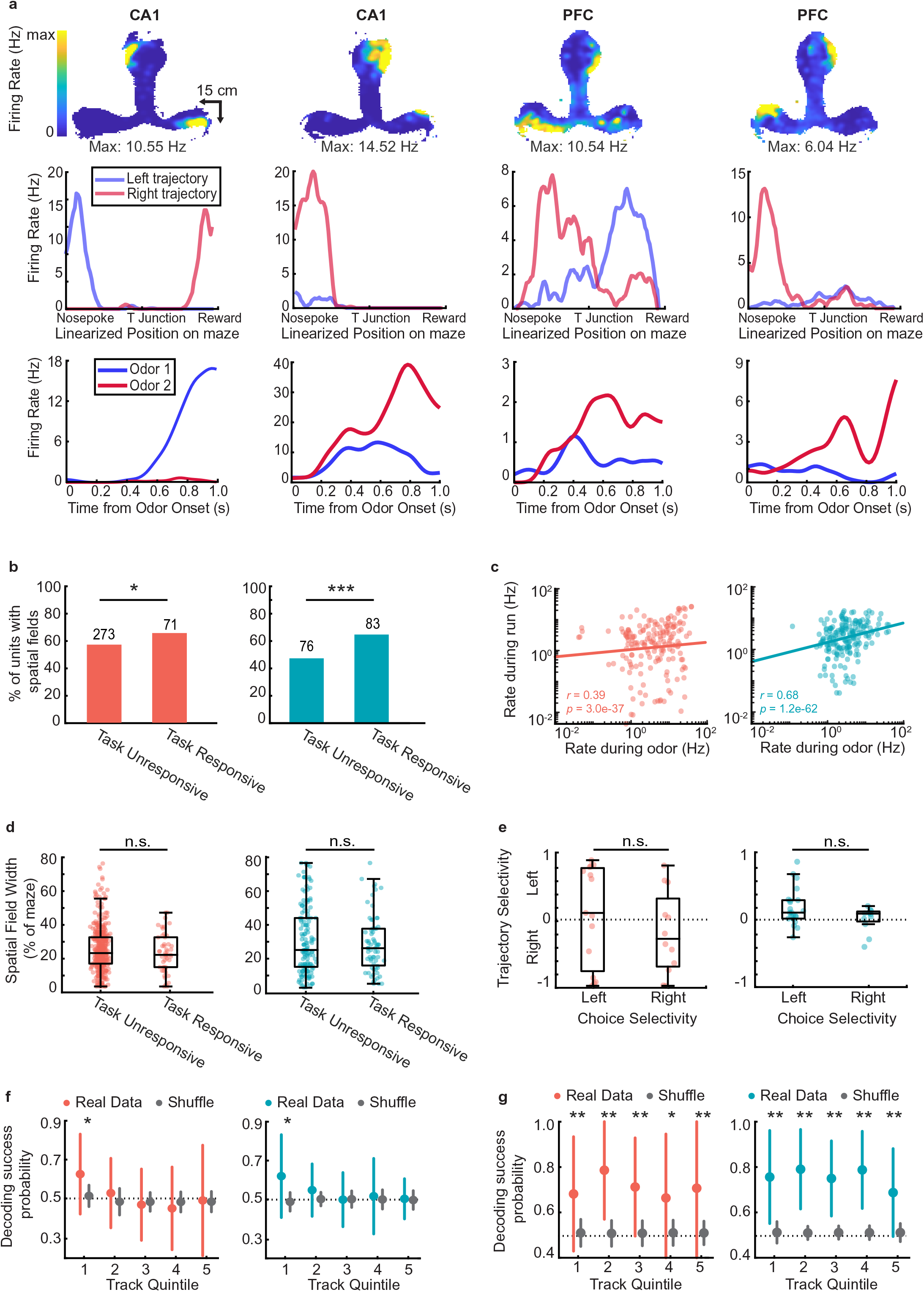
Representations of choice and space are maintained independently during delay period. a. Example choice-selective units (CA1, first two columns, PFC, last two columns) with spatial fields on the track. Top row: Heat map illustrating spatial fields during run bouts. Middle row: Linearized spatial tuning curves for outbound left and right run trajectories. Bottom row: PSTHs showing odor responses during decision-making period (Odor selectivity indices, left to right: 0.92, −0.43, −0.40, −0.57). b. Spatial field prevalence in task-unresponsive and task-responsive cells in CA1 (left) and PFC (right). CA1: 273/477 (57%) of task-unresponsive cells, and 71/108 (66%) of task-responsive cells had spatial fields (binomial test: p = 0.03*); PFC: 76/160 (48%) of task-unresponsive cells, and 83/128 (66%) of task-responsive cells had spatial fields (binomial test: p = 2.7e-5***). c. Correlation between firing rates during run and firing rates during odor sampling period for all putative pyramidal cells in CA1 (left) and PFC (right). (CA1: n = 585 cells, r = 0.39, p = 3.0e-37; PFC: n = 288 cells, r = 0.68, p = 1.2e-62). d. Spatial field width for odor unresponsive and odor responsive units in CA1 (left) and PFC (right). If a unit had a spatial field on both outbound trajectories, each field was counted separately. (Rank-sum tests; CA1: n = 452 fields, p = 0.15; PFC: n = 207 fields, p = 0.92). e. Trajectory selectivity scores of choice-selective units according to preferred choice (Rank sum tests, CA1: p = 0.26; PFC: p = 0.20). f. Decoding of choice identity by naïve Bayesian classifier according to CA1 (left) and PFC (right) task-responsive ensemble activity at five equally sized spatial quintiles along the full run trajectory of the maze. 1 spatial quintile = 24.6 cm (123 cm / 5). Colored error bars indicate real data mean ± s.d. whereas grey error bars indicate shuffled data mean ± s.d. Stars indicate quintiles where the fraction of correctly decoded trials was significantly higher than the fraction from the shuffled data (rank-sum tests, * = p < 0.05). g. As in (f), but decoding of choice identity according to CA1 (left) and PFC (right) spatially-modulated ensemble activity at each spatial quintile. (rank-sum tests, * = p < 0.05, ** = p < 0.01).

Cells that were task-responsive were more likely to have spatial fields on the track (odor sampling periods were excluded in spatial responses, see **Methods**), compared to cells that were not responsive during the odor sampling period (**Fig. 7b**, binomial test, CA1: p=0.029; PFC: p=2.7e-5). Additionally, for both CA1 and PFC neurons, we found that the firing rate during odor sampling was correlated with firing rate during running (**Fig. 7c,** CA1: r = 0.39, p = 3.0e-37; PFC: r = 0.68, p = 1.2e-62), and task-responsive cells had higher firing rates overall (**Supplementary Fig. 6b**), suggesting a possible relationship between cell activity during decision making and maintenance of the decision during the spatial delay. To examine this question further, we asked whether the spatial fields in either CA1 or PFC exhibited different characteristics based on whether they were responsive during the decision-making period. Surprisingly however, we found no difference in field width (**Fig. 7d,** rank-sum tests, CA1: p = 0.15; PFC: p = 0.92), field peak rate (**Supplementary Fig. 6c**), or field sparsity (**Supplementary Fig. 6d**) between cells that were task-responsive and task-unresponsive in either region.

Further, we also examined trajectory selectivity as animals ran through the central arm and on the side arms toward reward (examples of trajectory selective firing shown in **Fig. 7a,** second row). Trajectory selectivity was defined by comparing spatial tuning curves of cells on right versus left trajectories and calculating a selectivity index analogous to the choice selectivity index during the decision-making period (see **Methods**). Interestingly, although trajectory-selective cells were significantly more likely than trajectory-nonselective cells to respond in the decision-making period (**Supplementary Fig. 6e**), there was no relationship between the preferred trajectory during run periods and preferred choice during the odor-cued decision-making period (**Fig. 7e,** rank sum test CA1: p = 0.26; PFC: p = 0.20). Thus, although CA1 and PFC cells that were task responsive had higher firing rates and therefore higher engagement while traversing trajectories on the maze, there was no clear relationship between choice selectivity and trajectory selectivity in either CA1 or PFC.

To confirm this result, we sought to determine whether choice selective neuronal activity in the population during decision making persisted into the following run period. To examine this, the linearized T-maze was divided into 5 equally spaced quintiles and a Bayesian Classifier trained on odor period activity was used to decode the choice identity from the ensemble activity during run at each spatial quintile (see **Methods**; similar results were seen for spatial quartiles). We found that choice identity could be decoded at the first quintile of track, nearest to the odor port, but decoding accuracy diminished to chance level thereafter (**Fig. 7g**). In contrast, the animal’s upcoming behavioral choice could be accurately decoded at all periods along the trajectory if instead the spatially active ensembles in each respective spatial quintile was used as the training set (**Fig. 7h**). This suggests that the choice selectivity that emerges in ensembles during odor sampling (**Fig. 6**) persists transiently for a short period past the decision point, but that separate ensembles maintain choice-related information as animals traverse the spatial trajectory, possibly corresponding to theta oscillations during run (**Supplementary 2g**). This is further supported by the fact that, at least for CA1, choice-selective ensemble firing rate is highest within 20 cm of the odor port, and decreases thereafter (**Supplementary Fig. 6f**).These results together suggest that the choice-selective ensembles reflecting decision making during the odor-cue period transiently encode the choice during the initial track period, but the decision is subsequently maintained by a distinct ensemble during the working-memory period associated with the spatial delay on the central arm.

## DISCUSSION

These results provide new insight into the mechanisms of rhythmic coordination of hippocampal–prefrontal ensembles by beta rhythms for odor-cued associations and decision making. We found that during an odor-place associative memory task, beta and RR rhythms govern physiological coordination between the olfactory bulb, hippocampus, and PFC. Crucially, CA1-PFC beta coordination was uniquely elevated specifically during the decision-making portion of this task. During the odor sampling and decision-making period, task-responsive single unit and ensemble activity in the CA1-PFC network discriminates between choices and can reliably predict upcoming behavior. CA1-PFC beta coherence was elevated during the decision-making period, but we did not find evidence for difference in coherence for correct vs. incorrect trials. Instead, we found that phase-modulation of cells by beta rhythms was stronger on correct vs. incorrect trials, and in particular, CA1 pyramidal cells, as well as interneurons in both PFC and CA1 exhibited stronger phase modulation by these rhythms during accurate memory-guided decision making. This suggests that coordination by odor-driven beta rhythms may sculpt network activity through interneuron modulation, enabling emergence of task-related ensemble dynamics in local circuits to support decision making. Our results thus suggest that oscillatory modulation plays a key role in temporal evolution of ensemble dynamics in the CA1-PFC network for odor-cued decision making.

The mechanisms underlying encoding and recall of associations between sensory cues in the environment, and subsequent use of these associations to guide decisions, are of fundamental interest. Sensory cue-elicited recall and decision making during goal-directed behavior involves widespread networks encompassing sensory regions, medial temporal memory regions, and prefrontal executive function regions. Prominent odor-driven oscillations have been described in these areas (Frederick et al., 2016; Igarashi et al., 2014; Kay and Beshel, 2010; Kepecs et al., 2006; Lockmann et al., 2016; Nguyen Chi et al., 2016; Rangel et al., 2016; Stopfer et al., 2003; Tort et al., 2009), which can potentially coordinate these long-range networks to enable utilization of familiar olfactory cues to guide behavior. Here, the use of an odor-cued T-maze task allowed us to examine which rhythms enable coordination of olfactory-hippocampal-prefrontal networks, and whether these rhythms play a role in patterning ensemble activity in hippocampal-prefrontal network to enable memory-guided decision making.

In this task, the time period between odor onset and odor port disengagement provided a temporal window corresponding to odor-cued recall and priming of the subsequent decision to turn toward the reward location. We found prevalence of both beta (∼20-30 Hz) and RR (∼7-8 Hz) across the olfactory-hippocampal-prefrontal network during this odor sampling and decision-making period. The strength of beta coherence between the hippocampus and PFC was enhanced during the decision-making period compared to pre-odor periods as well as period of immobility during reward consumption. Further, while CA1-PFC RR coherence remained unchanged between odor-cued trials and trials where only air was presented, high beta coherence was specific to odor-cued trials corresponding to a decision-making task and was lower on air-cued trials corresponding to random choices. Based on these findings, we speculate that encountering a familiar odor stimulus and efficiently utilizing this associative memory for a decision, rather than respiration or movement preparation (Hermer-Vazquez et al., 2007), elicits the engagement of the hippocampal-prefrontal network by the beta rhythm.

Interestingly, although hippocampal SWRs have been linked to internal recall and planning (Buzsáki, 2015; Carr et al., 2011; Joo and Frank, 2018), there was very low prevalence of hippocampal SWRs during odor sampling, suggesting SWRs are not directly involved in odor-cued associative memory and decision making while the animal is re-encountering the familiar cue. Instead, our results indicate that beta rhythms play a key role in mediating this process, in addition to coordinating entorhinal-hippocampal networks for odor-cued recall, as described previously (Igarashi et al., 2014). In our experiments, rats were very familiar with the task and odor-place associations, with high performance levels. Therefore, the lack of SWRs during the decision-making period could reflect that these memories had been previously consolidated into neocortex during learning, resulting in a shift from largely hippocampal-dependent processing (Igarashi et al., 2014) to processing that is more reliant on hippocampal-cortical dialogue via the beta rhythm.

Odor-driven gamma rhythms at 40-100 Hz have also been reported in odor-memory tasks (Beshel et al., 2007; Frederick et al., 2016; Kay, 2014), and may play a complementary role with beta oscillations. It has been hypothesized that local olfactory networks may be governed by gamma rhythms for odor processing early in stimulus sampling, before shifting to a beta-dominant network state as downstream regions are engaged (Frederick et al., 2016; Martin et al., 2006). Our results corroborate this hypothesis by reinforcing the role of the beta rhythm in long-range coordination with a wider network outside the olfactory system for cognitive processing of odor stimuli. We speculate that sensory modality-specific rhythms may play a general role in cue-driven mnemonic decision making by coordinating sensory and cognitive areas, which can be tested in future studies.

Our analysis of single units in the hippocampus and prefrontal cortex revealed a group of cells that were selectively responsive in the odor task period, similar to previous reports in the hippocampus (Eichenbaum et al., 1987; Igarashi et al., 2014; Terada et al., 2017). However, we found that these selective cells often exhibited “opposite” selectivity on incorrect trials compared to correct trials, indicating that these cells do not respond solely to the perceptual qualities of the odor, in which case the response would be similar on all presentations of a particular odor regardless of the trial outcome. Instead, this response suggests coding of the behavioral choice associated with the odor cue, resembling findings from studies of combined cue modalities and cue-context associations (Allen et al., 2016; Ferbinteanu et al., 2011; Fitzgerald et al., 2011; Komorowski et al., 2009; McKenzie et al., 2014; Otto and Eichenbaum, 1992b; Schoenbaum and Eichenbaum, 1995b; Terada et al., 2017). Thus, the CA1-PFC neural response during odor sampling corresponds to the decision in response to the odor-place association. Future studies that combine recording from other olfactory regions, such as piriform cortex, together with hippocampus and PFC can potentially unveil how the cue representation evolves from odor perception to the recall-based decision.

Task-responsive neural populations exhibited differential activity patterns in response to the two odor-associations during decision making, and the animal’s upcoming choice could be predicted from this ensemble activity. Overall, the timing of ensemble odor discrimination preceded choice execution, indicating that the emergence of the neuronal representation can prime the cognitive decision. This emergence of neural ensembles that reflect associative memories governing decision making may also be indicative of a physiological mechanism for dynamic reactivation of memory engrams upon re-encountering the familiar odor stimuli (DeNardo et al., 2019; Liu et al., 2012). It remains unclear, however, how these neural dynamics are modulated by beta oscillations and how they evolve as animals learn new associations. One possibility is that they resemble the dynamics observed in the hippocampal-entorhinal circuit, in which coordination of the network at the beta rhythm and choice-selective neural activity emerge gradually as animals learn the task (Igarashi et al., 2014).

Similarly, CA1-OB beta coherence is strong during odor memory tasks and is known to increase over the course of learning (Martin et al., 2007). Although CA1-OB beta coherence was high during odor sampling in our task, we observed no significant relationship between CA1-OB coherence and task performance. Since previous reports suggest that this coherence is linked to the learning process, it is possible that a more direct relationship between coherence strength and performance would be observed only during learning, but not in this case where the odor association and task are familiar. Choice-coding ensemble dynamics may evolve in parallel with the increase in CA1-OB beta coherence over learning, similar to what has been reported in the hippocampal-entorhinal circuit (Igarashi et al., 2014). This possibility can be investigated by examining network dynamics and ensemble activity across multiple olfactory and cognitive areas during novel odor-place learning or reversal learning.

It is also noteworthy that we observed different mechanisms underlying odor-cued decision making versus subsequent maintenance of the representation of the choice during a spatial delay period. Odor-driven beta coherence and phase modulation has a role in accurate decisions, and possibly leads to emergence of ensemble selectivity which is predictive of the animal’s upcoming choices. However, these ensembles maintain their choice-selectivity only transiently after the decision execution as animals embark on the spatial trajectory toward reward. This selective coding is not maintained by the same ensembles during running, but instead other spatially modulated cells in CA1 and PFC maintain choice coding on the track, including the spatial delay period (common central arm on the T-maze) activity. Thus, beta driven ensembles mediate memory-guided decision making in the hippocampal-prefrontal network, but choice selective activity is then subsequently maintained by theta-driven spatially modulated activity, similar to reports in spatial working memory tasks (Taxidis et al., 2020).

The temporal evolution of ensemble activity underlying odor-cued decision making is reminiscent of temporal coding of odor stimuli mediated by oscillations in the olfactory system (Kepecs et al., 2006; Stopfer et al., 2003). Previous findings indicate that spike timing modulation according to the phase of prominent network rhythms can organize task-encoding neural ensembles to support memory guided behavior and decision-making (Benchenane et al., 2010; Buschman et al., 2012; Papale et al., 2016; Terada et al., 2017; Wikenheiser and Redish, 2015; Zielinski et al., 2019). Interestingly however, we found that although the strength of local beta phase-modulation was linked to correct decisions, putative neurons that encoded the choice were no more likely to be phase-locked than chance level. Further, we found evidence that different populations of cells are modulated by the two ongoing rhythms during decision making, implying that beta and RR represent two simultaneous modes of rhythmic coordination in the network (Rangel et al., 2016). This joint modulation of the underlying cell populations, along with our finding of strong cross-frequency coupling between the two rhythms, leads us to the interpretation that beta and RR coordinate the network cooperatively during odor-cued decision making. Previous reports of RR in olfactory processing (Karalis and Sirota, 2022; Kay, 2005; Kepecs et al., 2006; Nguyen Chi et al., 2016), along with evidence that multiple rhythms can simultaneously modulate ongoing processes (Lisman and Jensen, 2013; Rangel et al., 2016; Zhong et al., 2017), suggests that RR could be the dominant modulator coordinating the sensory element of the task, while beta coordination is key for employing the sensory cued association to make a decision. This interpretation is further supported by our results from un-cued air sessions, which did not involve memory-guided behavior, in which we found a reduction in beta coherence in the network but no change in RR coherence. This suggests that the sensory component of this process is maintained as the animal continues sniffing even in the absence of an explicit cue, whereas beta coordination is only engaged when a familiar cue is utilized for a decision.

We also found that a large population of interneurons were strongly modulated by the beta rhythm locally and the further that strength of this modulation was linked to task performance. PFC interneurons also showed temporal coordination with CA1 interneurons and pyramidal cells. This suggests that the beta rhythm can function by modulating interneuron activity, which sculpts ensemble dynamics in the local circuits governing decision making. In this manner, beta oscillations may play a role in establishing communication and organization of activity in sensory and cognitive networks enabling decisions based on cued associations. Thus, oscillatory phase-modulation may be indicative of a general network state that enables coordination.

## METHODS

All experimental procedures were approved by the Brandeis University Institutional Animal Care and Usage Committee (IACUC) and conformed to US National Institutes of Health guidelines. Eight male Long-Evans rats (3-6 months, 450-650 g) were used for experiments. Animals were housed individually in a dedicated climate-controlled animal facility on a 12-hour light/dark cycle. All experiments were carried out during light cycle. Upon arrival, animals were provided ad libitum access to food and water and handled regularly to habituate them to human contact.

### Behavior Apparatus

An olfactometer (MedAssociates Inc.) was used for dispensing odors. The olfactometer continuously dispensed clean air to the odor port until receiving a signal to open a solenoid valve which caused air to flow through liquid odorants, resulting in odorized air dispensed to the odor port. A vacuum tube attached to the odor port was used to continuously collect any residual odorized air between trials. Infrared beams were used at the odor port and reward wells to determine the precise timing of entry and exit from these areas.

### Odor-Place Association Task Training

Once rats reached a minimum threshold weight of 450 g, they were food restricted to no less than 85% of their free-feeding baseline weight. For initial behavioral training, rats were familiarized with the behavior room, the sleep box, running on a raised track to receive evaporated milk reward, and sniffing odors presented in the odor port. Following this habituation, rats were trained to hold their nose in the odor port for a minimum of 500 ms, with an auditory tone indicating when this 500-ms time threshold was reached. However, the rats could continue to sniff the odor for any longer duration of time, and odor would be continuously dispensed until they disengaged from the odor port. Throughout all training and experiments, if the rat exited the odor port before this threshold was reached, no reward was dispensed regardless of the rat’s choice and the rat was required to re-initiate the trial. These prematurely terminated trials were excluded from all analyses. For each trial, the odor sampling period was defined as the time from the onset when rat’s nose broke the infrared beam to offset when the beam break was terminated, and odor stimulus was stopped.

Animals were subsequently trained on the olfactory association. On each trial, one of two possible odors was dispensed at the odor port in a pseudo-random order. Odor 1 (Heptanol – pine/citrus scent) indicated that the rat should go to Reward 1 to receive reward. Odor 2 (ethyl butyrate – strawberry scent) indicated that the rat should choose Reward 2 (**Fig. 1a**). If the rat ultimately made the correct choice, evaporated milk reward would be dispensed at the chosen reward port upon triggering the infrared beam, whereas upon an incorrect choice no reward was dispensed.

Associative memory training was shaped in steps. The rats first learned the association between the odors and “right” vs “left” reward locations with reward wells close to the odor port. On subsequent training days, the reward wells were moved further and further away from the odor port, until rats could comfortably perform the task on the full T-maze (81 cm long center stem, and 43cm long reward arms). Two rats were not trained to run on the full T-maze, and instead were just required to perform the association with the reward wells within easy reach on either side of the odor port. For one animal, the maze had a shortened stem (40 cm instead of 81 cm), but the reward wells were still at the ends of the T arms. Task performance was calculated as the proportion of correct trials. Rats were trained until they could perform the task with at least 80% accuracy for 3 consecutive days.

Once training was complete, animals were once again provided with *ad libitum* access to food until they reached at least 600 g, before undergoing surgery. After surgery but before recording, rats were briefly re-trained on the association until they could again perform the task with at least 80% accuracy. On recording days, animals were allowed to continue performing the task until they reached satiation, about 100-150 trials per day. Task epochs were interleaved by sleep epochs, in which rats spent about 20 minutes in an opaque sleep box, with a sleep epoch as the first and last epoch of each day.

### Un-Cued Air Sessions

Four of the animals were tested on one session each in which only clean air was dispensed at the odor port, instead of two distinct odors. Reward was available at a randomly chosen reward port on each trial; although, since the trials were un-cued, the animals would often randomly choose the “incorrect” side and no reward was dispensed (**Supplementary Figure 3f**). All other aspects of the task were the same as the odor-cued task.

### Surgical Procedures

Surgical implantation techniques were performed as described previously (Shin et al., 2019; Tang et al., 2017; Tang et al., 2021). Animals were implanted with a microdrive array containing 30 independently moveable tetrodes targeting right dorsal hippocampal region CA1 (- 3.6 mm AP and +2.2 mm ML, 10-12 tetrodes), right PFC (+3.0 mm AP and +0.7 mm ML10-13 tetrodes), and right olfactory bulb (+7.2 mm AP and +0.8 mm ML, 1-4 tetrodes). Post-operative analgesic care was provided for 48 hours after surgery to minimize pain and discomfort. In two of the eight animals used for experiments, a nasal thermocouple was implanted in addition to the microdrive array to record the respiratory rhythm. The thermocouple was placed in the left nostril though a hole drilled in the skull at +7.5 mm anterior to the cribriform suture. The thermocouple wire was secured using dental acrylic and soldered to the same printed circuit board as the tetrodes.

### Tetrode Recordings

For 1-2 weeks following surgery, tetrodes were gradually lowered to the desired depths. Hippocampal tetrodes were targeted to the pyramidal layer of CA1 using characteristic EEG patterns (sharp wave polarity, theta modulation) and neural firing patterns as previously described (Jadhav et al., 2012; Jadhav et al., 2016). The final placement of tetrodes was confirmed in histological preparations using Nissl staining post-mortem. One tetrode in corpus callosum served as hippocampal reference, and another tetrode in overlying cortical regions with no spiking signal served as PFC reference. A ground screw (GND) installed in the skull overlying cerebellum also served as a reference, and LFP was recorded relative to this GND. All spiking activity was recorded relative to the local reference tetrode. Only LFP activity was recorded from the olfactory bulb. Electrodes were not moved at least 4 hours before and during the recording day to reduce drift, and were micro-adjusted at the end of each recording day to sample new cell populations.

Data were collected using a SpikeGadgets data acquisition system and software (SpikeGadgets LLC). Spike data were sampled at 30 kHz and bandpass filtered between 600 Hz and 6 kHz. LFP signals were sampled at 1.5 kHz and bandpass filtered between 0.5 Hz and 400 Hz. The animal’s position and running speed were recorded with an overhead color CCD camera (30 fps) and tracked by color LEDs affixed to the headstage.

### Data Analysis and Statistics

All data analysis was performed in MATLAB using custom code unless otherwise noted. Error bars indicate standard deviation unless otherwise noted. Significance was defined using an alpha of 0.05. Statistical details including tests used, p-values, and n values can be found in figure legends, and are described in-depth below.

### Local Field Potential

Trial-averaged spectrograms and coherograms were calculated using multi-taper spectral methods included in the Chronux package for MATLAB (www.chronux.org). To get the beta filtered local field potential (LFP) signal, raw LFP (with respect to GND) was band pass filtered at 20-30 Hz using a zero-phase IIR filter. Amplitude, phase, and envelope magnitude of the signals were obtained using a Hilbert transform. Beta power and coherence were z-scored to the epoch mean. Similarly, the respiratory rhythm signal was obtained by band pass filtering the raw LFP at 7-8 Hz.

### Bootstrap Tests

Bootstrap tests for **Supplementary Fig. 3e** were performed when coherence on odor-cued and air-cued trials were compared, to account for the much larger percentage of odor-cued trials. Data points on odor trials were randomly downsampled with replacement to match the number of air trials, and the mean was calculated for this new set of datapoints. This downsampling was done 1000 times, to create a new, bootstrapped distribution of means. The observed mean value for the air trial distribution was compared to the bootstrapped distribution. P-values were calculated by counting the number of values in the bootstrapped distribution that were greater than or equal to the observed mean value, and dividing by the total number of re-samples (1000). P-values less than 0.05 were considered statistically significant, and thus rejected the null hypothesis that the two observed distributions had equal means.

### Sharp-Wave Ripple Detection

Hippocampal sharp-wave ripples were detected as previously described (Jadhav et al., 2016; Shin et al., 2019; Tang et al., 2017). Briefly, the locally referenced LFP signal from CA1 tetrodes was filtered in the ripple band (150-250 Hz), and the envelope of the ripple-filtered LFPs was determined using a Hilbert transform. SWR events were detected as contiguous periods when the envelope stayed above 3 SD of the mean on at least one tetrode for at least 15 ms.

### Cross-frequency Coupling

Phase-amplitude coupling between RR and beta was computed for **Supplementary Fig. 2f** as previously described (Tort et al., 2010). In brief, the phases of RR were divided into 20 degree bins, and the mean amplitude of the beta rhythm at each phase was calculated. The mean amplitudes were then normalized by dividing each bin by the sum of amplitudes across all bins. The strength of phase-amplitude coupling was determined by comparing the amplitude distribution to a uniform distribution by calculating the modulation index (MI), which is based on the Kullback–Leibler (KL) distance but normalized so that values fall between 0 and 1, where a value of 0 indicates a uniform distribution of beta amplitudes across RR phases. Significance was determined by comparing the calculated MI to a null distribution generated by shuffling the trial number assignments of the RR phase series; this maintains the structure of the underlying rhythm but randomly aligns the beta amplitudes on each trial to each trial sequence RR phases.

### Single Unit Analysis

Spike sorting was done semi-automatically using MountainSort (Barnett et al., 2016; Chung et al., 2017), with manual curation. Only well-isolated units were used for analysis. Putative interneurons and pyramidal cells were classified based on average firing rate and spike width, as described previously (Shin et al., 2019; Tang et al., 2017). Units were classified as interneurons if they had an average firing rate exceeding 7 Hz and an average spike width under 0.3 ms. All other units were considered putative pyramidal cells. Units were excluded from analysis if they had fewer than 100 spikes across all sessions. A small number of pyramidal cells only had spikes during sleep epochs, and these were excluded from analysis (Jadhav et al., 2016; Karlsson and Frank, 2009) since all analysis done here focused exclusively on the task periods.

### Task Responsiveness

Task responsiveness was calculated as a change in firing rate following odor onset, compared to a pre-stimulus period of the same length of time as the odor sampling period on each trial leading up to odor port engagement. Sampling events were only considered if the animal held its nose in the odor port for longer than 0.50 seconds and proceeded to a reward port. Statistical significance was determined from the Wilcoxon signed rank test of those trial-matched rate differences for each odor identity separately. If a cell showed a significant change up or down from baseline for at least one of the odors, it was considered task responsive. Of note, there was only a single unit analyzed whose firing rate changed in opposite directions from baseline for the two odors, the remainder showed the same direction of change in firing rate for the two odors.

### Choice Selectivity

Firing rates during odor sampling were calculated as the number of spikes as a function of time from odor-port engagement to odor-port disengagement. Choice selectivity was calculated using the following equation:

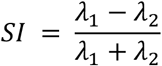

Where *λ*_1_ is the firing rate vector for Odor 1 trials, and *λ*_2_ is firing rate vector for Odor 2 trials. *SI* = 1 indicates that the cell only responded on Odor 1 trials, whereas *SI* = −1 indicates that the cell only responded on Odor 2 trials. To determine significance, a null distribution was generated in which the odor identities were shuffled across trials. Cells were considered choice-selective if the SI fell outside of 1.5 s.d. from the mean of the null distribution.

### Corrected Spike Cross- Correlogram

To correct for the triangular shape in the cross correlogram, we utilized a spike-shuffling procedure similar to that previously described (Kay et al., 2020). In brief, all spikes from one cell were jittered +/- 50 msec and the cross correlogram was recalculated from −.15 seconds to .15 seconds in 2.5 msec bins. This procedure was performed 1000 times, and the pointwise zscore of the real cross-correlogram was calculated. The highest peak within +/- 100 msec that achieved significance (p<.05), was chosen as the significant peak in the ccg, and only those cross correlograms in which there was a significant peak were displayed (and smoothed with a Gaussian kernel with σ=1 bin).

### Phase Locking

For all phase-locking analyses, the tetrode in each region with the most cells on each given day was used to measure oscillatory phase of either beta or RR. Phase locking of individual cells to the beta and respiratory rhythms was calculated by pooling the spike phases of each cell during the odor-sampling periods and performing a Rayleigh Z test for circular non-uniformity. When comparing phase coherence during correct trials versus that during incorrect trials, we adopted a down sampling strategy to adjust for rate differences, as described previously (Rangel et al., 2016). First, we matched the number of spikes during correct trials to that during incorrect trials. We then bootstrapped a mean vector length for the downsampled correct trials 1000 times and directly compared that to the MVL during incorrect trials. For calculation of the mean phase preference of each cell, we calculated the mean phase of all spikes during correct trials assuming a von-mises distribution.

### Principal Component Analysis (PCA)

We used PCA to visualize population activity of CA1 and PFC ensembles over time individual sessions. Spiking activity in the decision making period aligned to odor onset was binned (binsize = 100 ms, window = 0-1 s), and a firing rate matrix was constructed where each row represents a bin and each column represents a neuron. Only sessions with at least 4 task-responsive neurons (threshold applied for both CA1 and PFC; CA1 and PFC ensembles were examined separately) were used for analyses. We used PCA to find the principal component coefficients of the matrix, and applied the coefficients to the population activity for left-bond vs. right-bound trials. Population activity was projected onto the PC space. The first 3 PCs were used for visualization and analysis. The Euclidean distance between left-bound and right-bound average trajectories was compared to a chance-level distance distribution computed by shuffling the trial identities across trials and creating a null distribution. The timepoint at which the population responses were considered significantly distinct from each other was defined as the first timepoint at which the real distance between trajectories surpassed the 95% confidence interval of the null (shuffled) distribution. Similar results were obtained for Euclidean distance computed with the first 3 PCs and for Euclidean distance computed for all neurons without dimensionality reduction.

### GLM

A generalized linear model (GLM) with a log link function was constructed to predict reward choice based on neural activity during the odor sampling period. Activity from all task-responsive neurons that were active on at least 10 trials was included. Neural population activity from different length time bins from 0.1 seconds to 1 second aligned to odor onset was used for prediction, i.e. 0-0.1s, 0-0.2s … 0-1.0s. Five-fold cross validation was used to test prediction. For each fold, the session’s trials were randomly partitioned into five equally sized sets. Four of the five sets were used to train the GLM model and the remaining set was used to test. The prediction accuracy was calculated by dividing the number of correctly predicted trials by the total number of trials used for testing for each fold.

Significance was determined by performing the same procedure as described above but shuffling the trial outcomes to obtain a null distribution. At each time bin used for the prediction, a rank-sum test was performed on the prediction accuracy using the real data compared to the shuffled data.

### Occupancy Maps

For all spatial field analyses, only the 5 animals that were trained on the full T-maze were used; the three animals trained on the truncated maze were excluded due to insufficient spatial data. Two-dimensional occupancy maps were generated by calculating occupancy in 2-cm square spatial bins from epochs in which the animals running speed exceeded 3 cm/sec and convolving with a 2d Gaussian (**σ**= 2 pixels).

### Place Field Determination

2-D occupancy-normalized firing rate maps were generated by dividing the spikes at each 2-D pixel by the unsmoothed occupancy at that pixel, and then smoothing with a Gaussian kernel of (σ= 2 pixels, or 4 cm).

Linearized trajectory occupancy maps were calculated as previously described^31,97,98^. Briefly, the rats 2d spatial coordinates were first segmented into run epochs based on contiguous bouts of time in which the animals running speed exceeded 3 cm/sec. Then, the position of the animal during each ‘run epoch’ was categorized by its origin and destination, and each position was assigned to one of 100 (1.23 cm) bins from the origin to the destination of that route. Then the total number of spikes at each bin was divided by that bin’s occupancy, and the map was convolved by a Gaussian kernel with a standard deviation of 2 pixels (4 cm).

Place field peak was calculated as the peak rate bin on the smoothed, linearized rate map, and place fields were only considered for trajectories in which the cell had a peak firing rate of at least 1 Hz. Place field width was calculated using a flood fill-algorithm in which the edges were defined as the closest bins to the peak bin in which the rate fell below 25% of the peak rate. Place field sparsity and information scores were calculated as previously described (Skaggs et al., 1993). A cell was determined to have a place field if its peak firing rate along that trajectory was ≥ 2 S.D. above the mean of a bootstrapped distribution generated by circularly shifting the spikes in time for each individual run, and if the field covered less than 75% of the linearized trajectory. When analyzing the place field characteristics of choice-selective cells (**Fig. 7**, **Supplementary Fig. 6**), the field of each outbound journey was analyzed separately so as to prevent ‘choosing’ certain fields over others.

### Trajectory Selectivity

Trajectory selective cells were identified using a previously validated method (Shin et al., 2019). Briefly, the linearized spatial tuning curve was calculated separately for each outbound trajectory and the correlation between those trajectories was computed. A cell was identified as trajectory selective if this correlation value exceeded the 95^th^ percentile of a distribution wherein each outbound trajectory identity was shuffled and the correlation recalculated.

To measure trajectory selectivity as it related to decision period selectivity, we used an analogous method to the odor selectivity index. Briefly, we calculated the mean firing rate along each run beginning one half second following odor port exit and once the animal’s speed exceeded 3 cm/sec to the end of that run (when the animal either reached the goal or its velocity fell below 3 cm/sec for more than ½ second). The difference in these mean firing rates across runs was divided by the sum of those two mean rates to generate a trajectory selectivity index.

### Bayesian Decoder

Choice identity decoding was performed as previously described (Karlsson and Frank, 2009). Briefly, a memoryless Bayesian decoder was built for each of the two choice identities from spikes occurring in the odor sampling period. Then, the likelihood of each choice (x) was reconstructed from their posterior probabilities given the spikes occurring at each segment along the maze on each run (p(x | spikes) = p(spikes | x) * p(x) / p(spikes)). Run activity was defined as contiguous bouts following the odor sampling period when the animal was traveling above 3 cm/sec. Additionally, to prevent overlap between odor-period spiking and run spiking, the run periods were defined beginning 0.5 seconds following odor port disengagement time. We assumed that the N active cells fired independently and followed a Poisson process, giving the following equation.

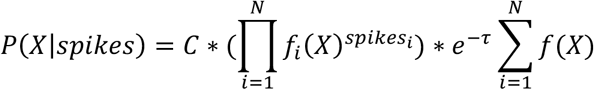

Where C is a normalization constant across the two choice identities. The p-value was calculated from the Gaussian statistics of a Monte Carlo random shuffle (200 shuffles) of choice identity during the odor sampling period. When choice identity decoding was calculated from the likelihoods of activity from each run, the decoding was performed in the same 5-fold leave-one-out fashion as was used in the GLM analyses, and the p value was calculated from the same Monte Carlo random shuffle of the training set route identities as the odor period based decoding.

### Data Availability

The behavioral and electrophysiological data supporting the findings in this study are archived on servers at Brandeis University and are available upon request to the corresponding author.

### Code Availability

All data processing, analyses and statistics in this study were conducted using open-source package Mountainsort (https://github.com/flatironinstitute/mountainsort) and custom code in MATLAB (R2018), unless otherwise noted. All custom code is available on GitHub and upon request to the corresponding author.

## ACKNOWLEDGMENTS

This work was supported by NIH Grant R01MH120228, and a Smith Foundation Odyssey award to S.P.J. We thank the late Howard Eichenbaum for assistance with the behavioral task.

## AUTHOR CONTRIBUTIONS

SPJ: Conception and Study Design, Data Analysis; CAS: Experiments, Data Analysis, JHB: Data Analysis, EK: Experiments, PM: Data Analysis. SPJ, CAS and JHB wrote the manuscript with input from all authors.

## COMPETING INTERESTS STATEMENT

The authors declare no competing interests.

---

**Supplemental Figure 1 (Related to Figure 1).**
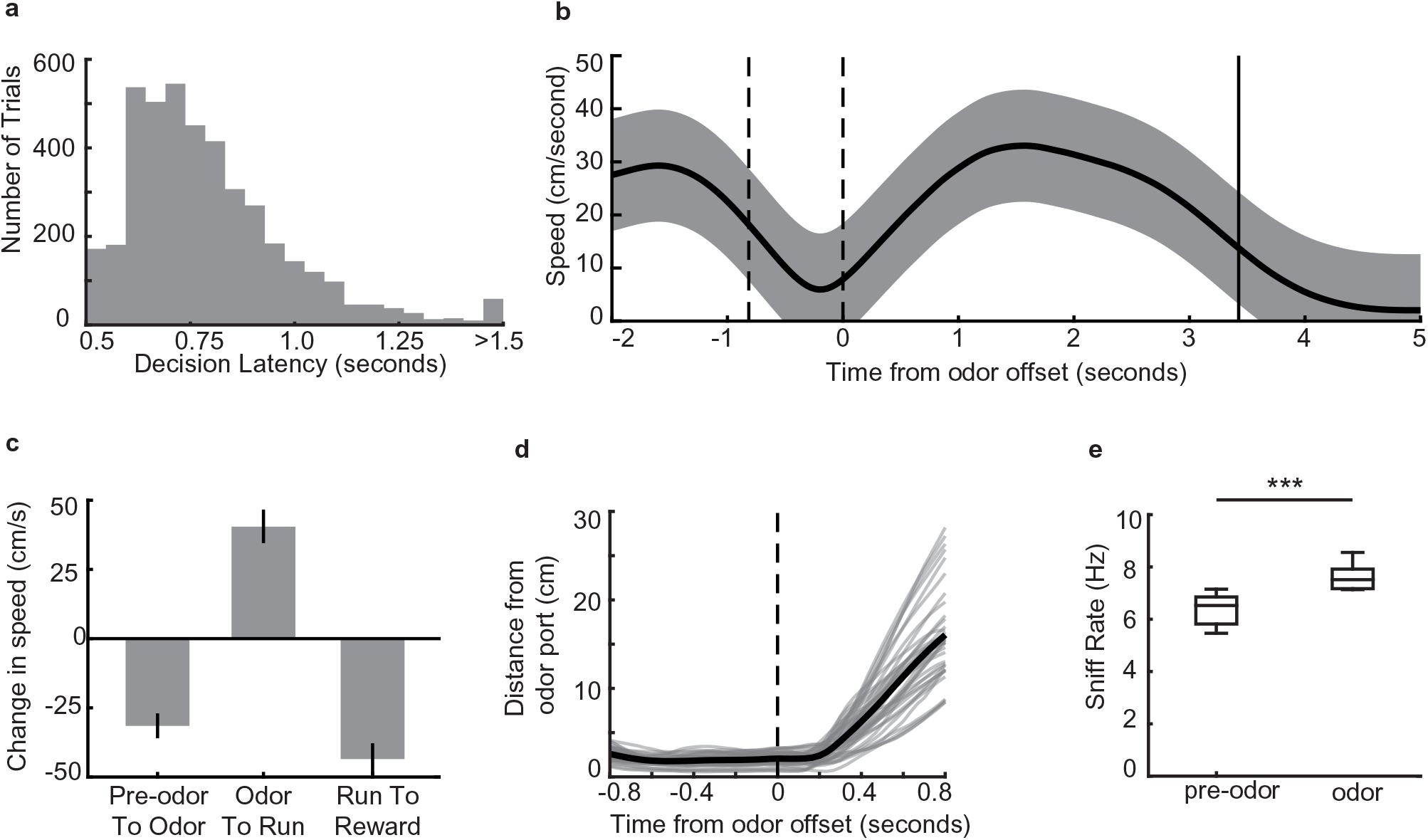
Behavioral parameters on the odor-place associative memory and decision-making task. a. Histogram of decision latencies. Mean: 0.82 seconds ± 0.25 s.d. b. Mean run velocity across the full task for an example session from one animal, aligned to odor offset. Area between dashed lines indicates average odor-sampling period, solid line indicates average reward onset time. Shaded area indicates s.d. c. Average change in speed across different task epochs, for all animals that ran on the full maze (and not the truncated maze) (n = 5). Error bars indicate s.d. d. Distance from odor port over time aligned to odor port offset time. Grey lines are individual sessions, black line is mean across sessions. (n = 38 sessions). e. Sniff rate as measured by the thermocouple signal, during odor (7.1 ± 0.39 Hz, mean ± s.e.m.) and time matched pre-odor periods (6.2 ± 0.29 Hz, mean ± s.e.m.). Box plots indicate interquartile ranges (signed-rank test, n = 11 sessions across 2 animals, p = 9.8e-4***).

**Supplemental Figure 2 (Related to Figure 2).**
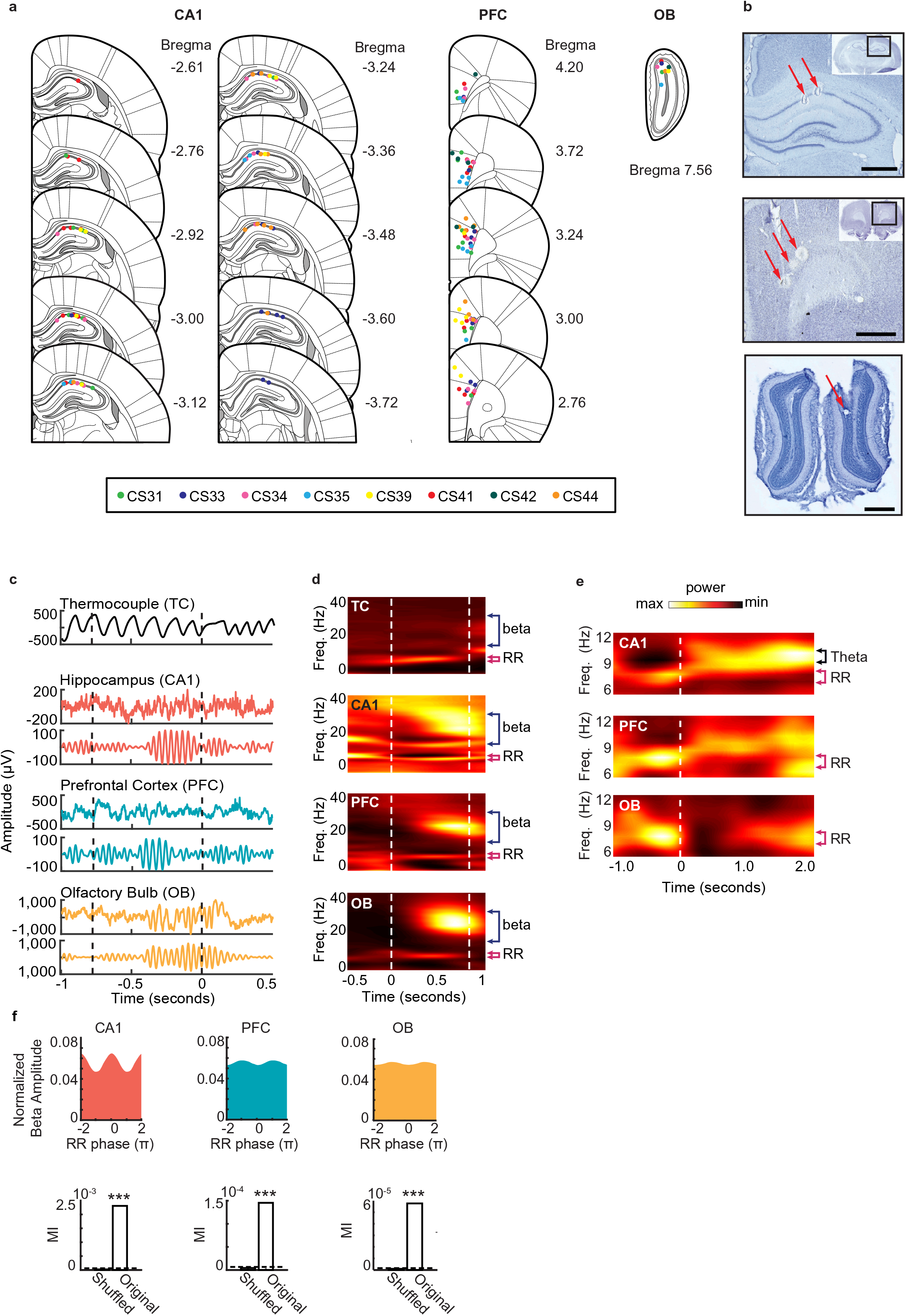
Rhythmic activity in the olfactory-hippocampal-prefrontal network. a. Recording locations for 8 rats (color coded) where tetrode locations were recovered. The recording sites were reconstructed from the electrolytic lesions in *post hoc* Nissl-stained coronal brain sections and mapped onto the stereotaxic atlas^1^. Electrodes were localized to target areas in dorsal area CA1, medial prefrontal cortex (PFC), and granule cell layer of OB after histology (PFC electrodes localized primarily to prelimbic cortex, with a few electrodes in anterior cingulate cortex (ACC)). b. Sample Nissl-stained brain sections showing final tetrode placement in hippocampus (top), PFC (middle), and olfactory bulb (bottom). Arrows indicate tetrode lesions. Scale bars each represent 1 mm. c. Examples of thermocouple and LFP traces from one tetrode in each region during presentation of odor from one trial, aligned to odor port disengagement. Area between dashed lines indicates odor sampling period. Top to bottom: Respiratory rhythm recorded via thermocouple, CA1 signal, beta band (20-30 Hz) filtered CA1 signal, PFC signal, beta band filtered PFC signal, OB signal, beta band filtered OB signal. d. Time-frequency plots showing power spectra across all animals aligned to odor onset (at time 0). Color scale represents z-scored power. Area between dashed lines indicates average odor sampling period. Beta band is marked by blue bracket, and RR band is marked by pink bracket. Thermocouple signal (TC), n = 12 sessions, max 0.92, min −0.30; CA1: n = 38 sessions, max 0.29, min −0.33; PFC: n = 38 sessions, max 0.58, min −0.15; and OB: n = 38 sessions, max 2.28, min −0.20. e. Time-frequency spectrograms across animals that ran on the full maze, aligned to odor port disengagement (at time 0) and extending into the run period on the track. Color scale represents z-scored power for each region: CA1: max 0.45; min −0.34, PFC: max 0.21; min −0.14, and OB: max 0.72; min −0.16. Dashed line indicates odor port disengagement time. RR band is marked by pink bracket. f. Phase-amplitude coupling between RR and beta, in CA1 (left), PFC (middle), and OB (right). Top row: normalized phase-amplitude histograms. Bottom row: modulation index (MI)^2^ of real data compared to the MI from a trial-shuffled dataset (500 shuffles). Dashed line indicates significance level (alpha = 0.05, p < 0.001***).

**Supplemental Figure 3 (Related to Figure 2).**
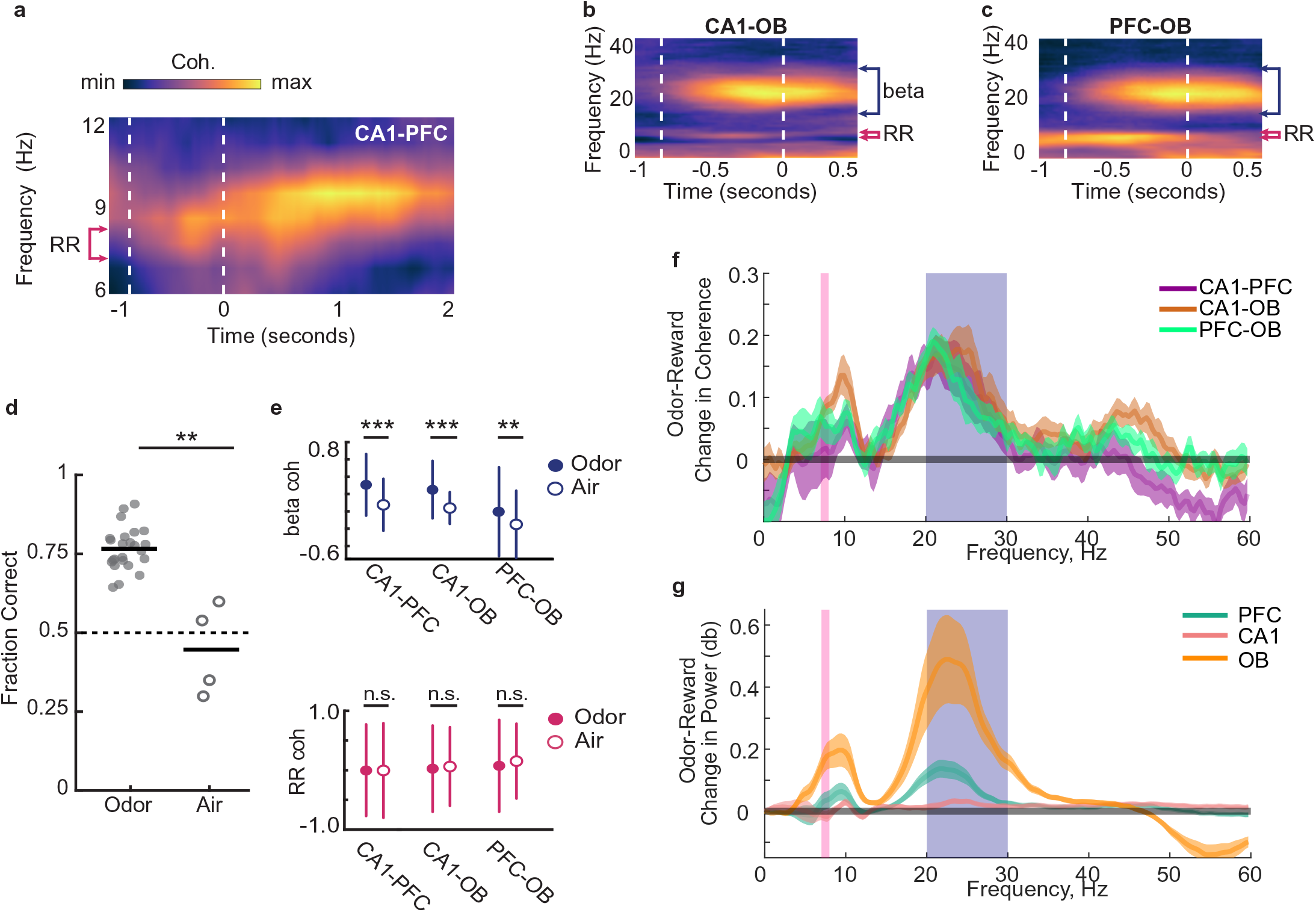
Beta coordination underlies decision making based on odor-place associations. a. CA1-PFC coherence across animals that ran on the full maze, aligned to odor port disengagement and extending into the run period on the track. Color scale represents z-scored coherence. Min = −0.18, max = 0.50. Dashed line indicates odor port disengagement time. RR band is marked by pink bracket. b. Top: CA1-OB coherence spectra across all animals and all tetrode pairs (n = 38 sessions). Area between vertical dashed lines indicates mean decision-making period, aligned to odor port disengagement. Color scale represents z-scored coherence (max 0.41, min −0.25). Middle: Beta (15-30 Hz) coherence during odor periods and time-matched pre-odor periods. (signed-rank test, n = 38 sessions, p = 1.7e-7***). Bottom: RR (7-8 Hz) coherence during odor periods and time-matched pre-odor periods. (signed-rank test, p = 6.3e-4***). c. As in (b), but for PFC-OB coherence. Top: max 0.45, min −0.28. Middle: signed-rank test, p = 4.3e-7. Bottom: signed-rank test, p = 0.30. d. Performance of three animals on the odor-cued task (n = 24 sessions) versus the air-cued task (n = 4 sessions) in which animals were presented with only air as a neutral stimulus instead of two distinct odors. Black bars indicate mean. Dashed line indicates chance level. (Rank-sum test, p = 0.002**) e. Beta (top row) and RR (bottom row) Z-scored coherence between CA1-PFC, CA1-OB and PFC-OB on correct trials with odors (n = 3174) and randomly rewarded trials with only air presented at the odor port (n = 134). Correct trials were randomly subsampled with replacement to match the number of incorrect trials 1000 times. Error bars indicate s.d. of real data distributions. (Bootstrap tests: beta: CA1-PFC: p < 0.001***, CA1-OB, p < 0.001***, PFC-OB: p = 0.002**; RR: CA1-PFC: p = 0.49, CA1-OB, p = 0.30, PFC-OB, p = 0.12) f. Change in coherence from reward period to odor period. Beta coherence during odor sampling was higher for all pairs (sign rank test; CA1-PFC p-0.01, CA1-OB 0=0.0012, PFC-OB p=0.0024) and RR coherence was higher only for OB-CA1 and OB-PFC (CA1-PFC p=0.56, CA1-OB p=0.0044, PFC-OB p=0.64e-4) g. Change in power from reward period to odor period. Beta power increased in all three regions from reward to odor sampling (sign rank test, CA1 p=0.0012, PFC p=5.3e-4, OB p=4.4e-4) whereas RR power only increased in OB (CA1 p=0.062, PFC p=0.11, OB p=4.4e-4)

**Supplemental Figure 4 (Related to Figures 3 and 4).**
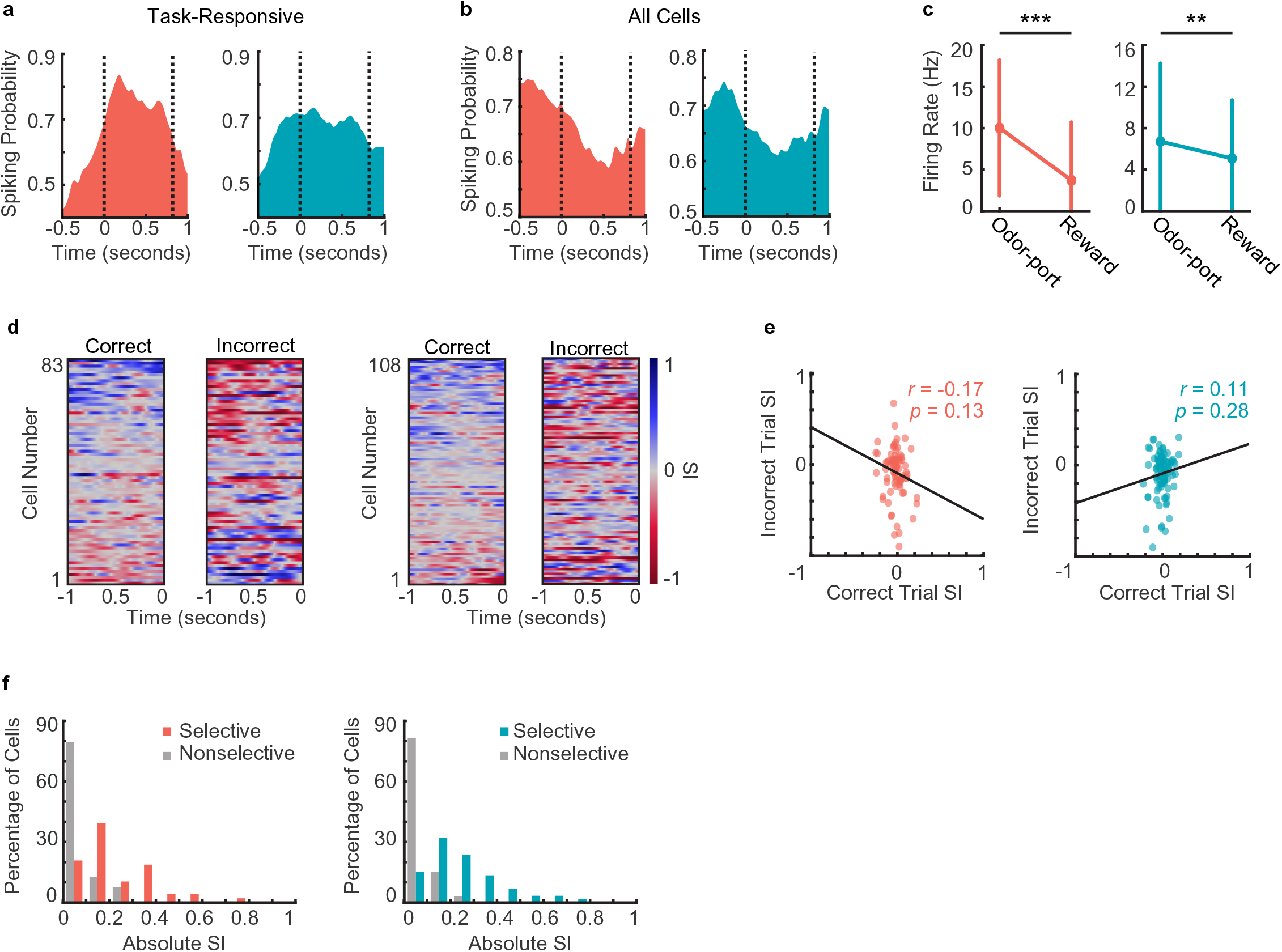
Neural responses during decision-making period. a. Spiking probability during decision-making period for all task-responsive cells in CA1 (left, n = 125 cells) and PFC (right, n = 157 cells). Area between the dashed lines represents decision-making period. b. Spiking probability during decision-making period for all cells (task-responsive and task unresponsive, putative pyramidal cells and interneurons combined) in CA1 (left, n = 917 cells) and PFC (right, n = 507 cells). Area between the dashed lines represents decision-making period. c. Firing rates of task-responsive pyramidal neurons during odor sampling and reward consumption (Signed-rank tests. CA1 (left): n = 125 cells, p = 5.1e-13; PFC (right): n = 157 cells, p = 0.009). d. Selectivity index (SI) of all task-responsive but not choice-selective pyramidal cells in CA1 (left 2 panels) and PFC (right 2 panels) on correct trials and incorrect trials, aligned to odor port disengagement. Cells are sorted according to peak selectivity on correct trials and sorting order is the same for both plots. e. Correlation between correct trial SI and incorrect trial SI for all non-choice-selective pyramidal neurons. (CA1 (left): n = 83 cells, r = −0.34, p = 0.001**; PFC (right): n = 108 cells, r = −0.34, p = 3.0e-4***). f. Histograms showing absolute selectivity indices for choice selective vs. non-choice-selective pyramidal cell populations in CA1 (left) and PFC (right).

**Supplemental Figure 5 (Related to Figure 5).**
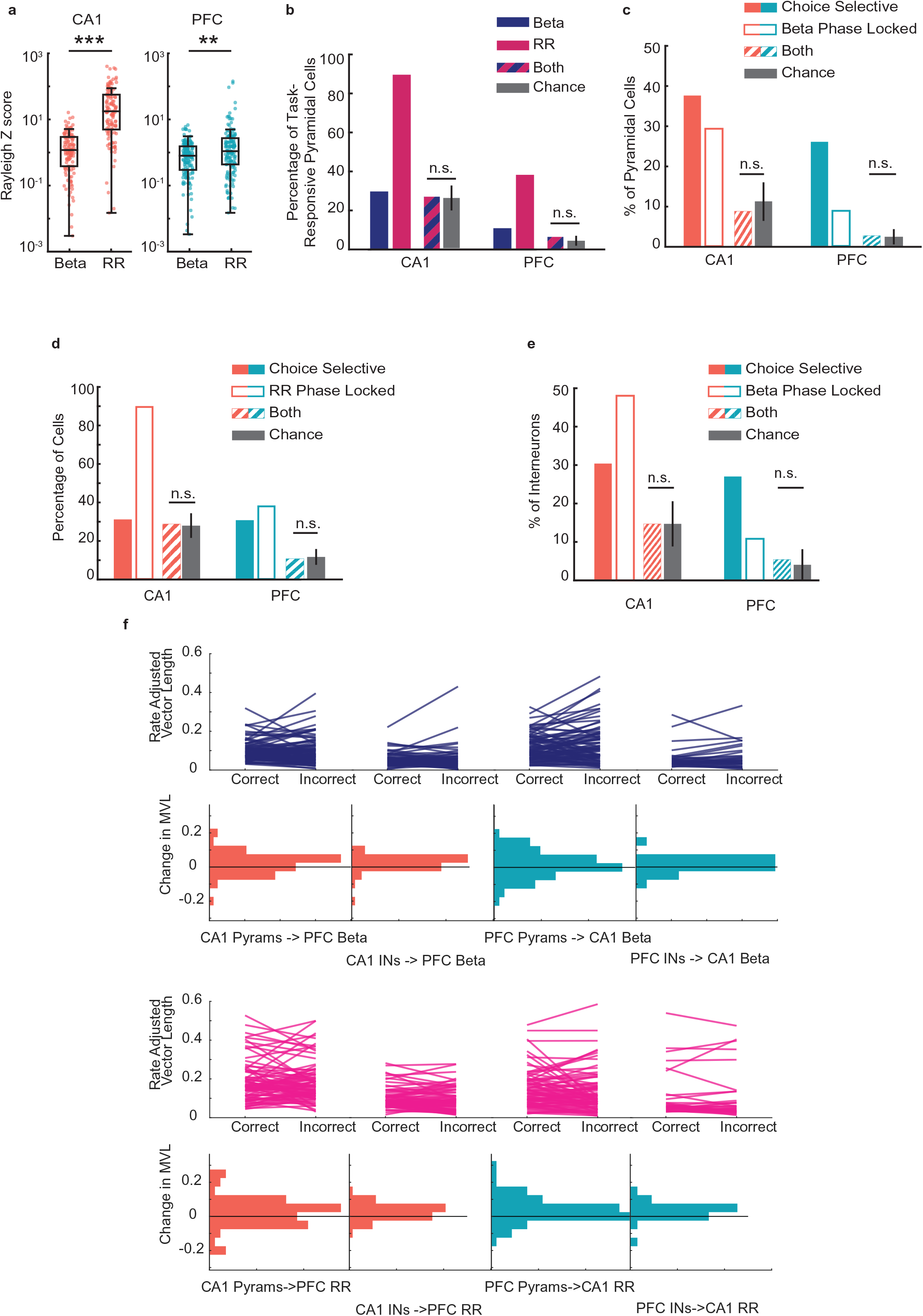
Phase locking to respiratory rhythm. a. Rayleigh Z scores for spike-phases of task-responsive cells for beta and RR (rank-sum test; CA1: n = 125 cells, p = 3.2e-26***; PFC: n = 157 cells, p = 0.002**). b. Percentage of task-responsive pyramidal cells in CA1 and PFC that were phase locked to beta, RR, or both beta and RR. Grey bars indicate chance level of cells being phase locked to both rhythms, error bars indicate the range of cell percentages that fall within the 95% bounds of the chance binomial distribution. Binomial tests. CA1: p = 0.08; PFC: p = 0.06. c. Percentage of cells that are choice-selective (solid bars), phase-locked to the beta rhythm (outlined bars), and both choice-selective and phase-locked (striped bars). Grey bars indicate chance level of cells being both choice-selective and phase-locked, error bars indicate the range of cell percentages that fall within the 95% bounds of the chance binomial distribution. (binomial test, CA1: p = 0.08; PFC: p = 0.16). d. Percentage of cells that are choice-selective (solid bars), phase-locked to RR (outlined bars), and both choice-selective and phase-locked (striped bars). Grey bars indicate chance level of cells being both choice-selective and phase-locked, error bars indicate the range of cell percentages that fall within the 95% bounds of the chance binomial distribution (Binomial test, CA1: p = 0.077; PFC: p = 0.10). e. As in (c), but for interneurons. (binomial test; CA1: p = 0.11; PFC: p = 0.20) f. Rate adjusted vector length (top) and histogram of change in vector length from correct to incorrect trials (bottom) for all task responsive neurons in PFC and CA1 relative to cross-region Beta and RR. No comparisons revealed significant results. (signed-rank tests, all p>.05)

**Supplemental Figure 6 (Related to Figure 7).**
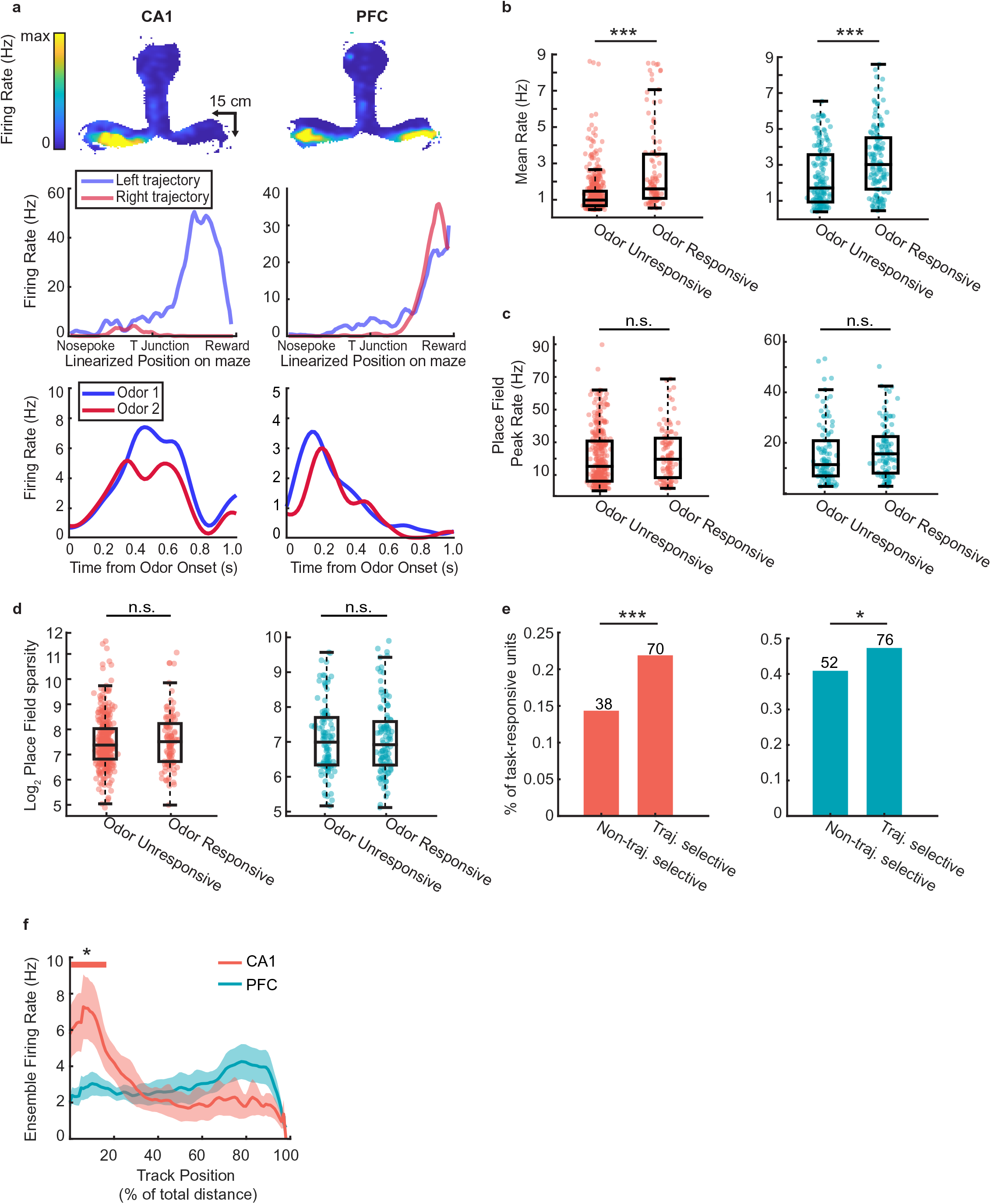
Spatial activity of task-responsive and task-unresponsive neurons. a. Example task-responsive (but not choice-selective) units from CA1 (left) and PFC (right) with spatial fields on the track. Top row: Heat map of firing fields during run bouts. Middle row: Linearized spatial tuning curves for outbound left and right run trajectories. Bottom row: PSTHs showing odor responses during decision-making period. b. Mean firing rate of task-responsive and task-unresponsive units during running on the maze, excluding decision-making periods (Rank-sum tests: CA1 (left): n = 585 cells, p = 2.8e-14***; PFC (right): n = 288 cells, p = 1.3e-5***). c. Place field peak rate for task-responsive and task-unresponsive cells. (Rank sum tests, CA1 (left): n = 452 fields, p = 0.14; PFC (right): n = 207 fields, p = 0.08). d. Place field sparsity for task-responsive and task-unresponsive cells. (Rank sum tests, CA1 (left): p = 0.70; PFC (right): p = 0.88). e. Percentage of trajectory selective and non-trajectory selective cells that that were also task-responsive (Binomial tests: CA1 (left): n = 108 odor responsive units, p = 3.0e-4***; PFC (right): n = 128 odor responsive units, p = 0.02*). f. Mean ensemble firing rate of task-responsive units in CA1 and PFC during outbound runs. Solid line indicates mean, shaded area represents s.d. Thick colored bar above marks positions in which the ensemble rate exceeds the 99th percentile of a bootstrap randomized null mean ensemble rate.

## Notes

### Competing Interest Statement

The authors have declared no competing interest.

